# Preservation of prehearing spontaneous activity enables early auditory system development in deaf mice

**DOI:** 10.1101/2022.11.01.514787

**Authors:** Calvin J. Kersbergen, Travis A. Babola, Patrick O. Kanold, Dwight E. Bergles

## Abstract

Intrinsically generated neural activity propagates through the developing auditory system to promote maturation and refinement of sound processing circuits prior to hearing onset. This early patterned activity is induced by non-sensory supporting cells in the organ of Corti, which are highly interconnected through gap junctions containing connexin 26 (*Gjb2*). Although loss of function mutations in *Gjb2* impair cochlear development and are the most common cause of congenital deafness, it is not known if these mutations disrupt spontaneous activity and the developmental trajectory of sound processing circuits in the brain. Here, we show in a new mouse model of *Gjb2-*mediated congenital deafness that cochlear supporting cells unexpectedly retained intercellular coupling and the capacity to generate spontaneous activity, exhibiting only modest deficits prior to hearing onset. This coordinated activation of IHCs led to coincident bursts of activity in central auditory neurons that will later process similar frequencies of sound. Despite alterations in the structure of the sensory epithelium, hair cells within the cochlea of *Gjb2* deficient mice were intact and central auditory neurons could be activated within appropriate tonotopic domains by loud sounds at hearing onset, indicating that early maturation and refinement of auditory circuits was preserved. Only after cessation of spontaneous activity following hearing onset did progressive hair cell degeneration and enhanced auditory neuron excitability manifest. This preservation of cochlear spontaneous activity in the absence of connexin 26 may increase the effectiveness of early therapeutic interventions to restore hearing.

## INTRODUCTION

Non-sensory supporting cells in the cochlea comprise a molecularly and physiologically diverse group of cells required for hearing. They secrete trophic factors critical for synapse development, remove neurotransmitters and ions generated through sound-evoked mechanotransduction, and provide structural support to the sensory epithelium to enable amplification^1, 2^. Despite their remarkable diversity, cochlear supporting cells are extensively coupled to each another through gap junctions, forming a functional syncytium to coordinate development of this complex structure and allow ionic and metabolic homeostasis to sustain hearing^3–7^. These intercellular channels consist primarily of the gap junction protein connexin 26 (Cx26) encoded by *Gjb2*. Although not directly involved in mechanotransduction or signal transmission from hair cells to spiral ganglion neurons (SGNs), mutations in *Gjb2* are the most prevalent cause of congenital non-syndromic hearing loss, accounting for >25% of all genetic hearing loss worldwide^8–10^. Prior studies revealed that loss of function variants in *Gjb2* impair the structural development of the cochlea and result in deafness^11–15^. Nevertheless, cochlear implants can restore hearing in children that lack Cx26^16^, suggesting that maturation of central auditory pathways occurs despite disruption of normal cochlear function during a critical period of development. However, the mechanisms that enable early maturation of brain circuits that process acoustic information despite these cochlear abnormalities have not been defined.

All sensory pathways that have been examined experience a period of early refinement dependent on input from peripheral sensory organs^17^. In the auditory system, spontaneous bursts of action potentials emerge from the cochlea prior to hearing onset (time of ear canal opening when there is a profound increase in sensitivity to airborne sound) and propagate throughout the developing central auditory pathway^18–23^, inducing correlated firing of neurons that will later process similar frequencies of sound. This intrinsically generated activity is initiated by a group of inner supporting cells (ISCs) within the sensory epithelium that reside adjacent to inner hair cells (IHCs). Stochastic, spontaneous release of ATP from individual ISCs triggers a cascade of events that culminates in depolarization of nearby IHCs, producing highly stereotyped activity in the absence of mechanosensation. In mouse, this rhythmic burst firing continues for almost two weeks after birth, allowing auditory neurons to engage in experience-dependent plasticity to ensure proper processing of peripheral stimuli when the ear canal opens and acoustic sensitivity increases dramatically^17, 24, 25^. Disruption of Cx26-mediated communication during this period alters developmental programs in the cochlea, preventing formation of fluid filled spaces around hair cells (e.g. Tunnel of Corti and Space of Nuel)^11, 12^. It may also impair some forms of extracellular communication, as Cx26 can exist as an unpaired hemichannel at the cell surface to enable release of diverse signaling molecules, including ATP^26, 27^. However, the effect of Cx26 deficiency on spontaneous activity has not been examined, and the consequences for maturation and refinement of auditory processing circuits within the brain remains poorly understood.

To understand how Cx26 deficiency influences both spontaneous activity in the cochlea and neuronal activity patterns in central auditory centers, we developed a new mouse model to achieve targeted, constitutive deletion of Cx26 (Cx26 cKO mice) from pre-hearing supporting cells within the organ of Corti, as global Cx26 deficiency is lethal^28^. Cx26 cKO mice exhibited a profound loss of acoustic sensitivity at ear canal opening (P11) and progressive cellular degeneration within the cochlea, consistent with evidence that supporting cell coupling within the cochlea is required for normal hearing. However, ISCs in cochleae from prehearing Cx26 cKO mice unexpectedly retained low membrane resistance, intercellular dye coupling, and spontaneous electrical activity mediated by ATP release. These ISCs continued to induce coordinated activation of IHCs along the length of the cochlea, excitation of SGNs, and correlated firing of auditory neurons within isofrequency zones *in vivo*. Although auditory neurons in the midbrain and cortex after ear canal opening were only activated in Cx26 cKO mice by very loud sounds, their spatial and temporal neuronal activity patterns were remarkably similar to controls. With increasing age, cortical neuron responses to sound in Cx26 cKO mice became larger and more prolonged, suggesting an adaptive response to the lack of input after cessation of spontaneous activity. These results reveal that cochlear spontaneous activity is preserved in the absence of Cx26, enabling activity dependent maturation of auditory circuits and homeostatic control of acoustic sensitivity despite later deafness, helping to establish neural networks in the brain that can be engaged by cochlear prostheses.

## Results

### Targeted deletion of Cx26 from the cochlear sensory epithelium leads to auditory dysfunction

Non-sensory supporting cells in the organ of Corti maintain electrical and chemical coupling through gap junctions, which have been implicated in generating spontaneous activity prior to hearing onset by enabling transfer of second messengers between cells^29–31^ and ATP release when present as unpaired hemichannels at the cell surface^18, 26^. To determine how selective loss of Cx26 from these supporting cells influences spontaneous activity before hearing onset and later auditory function, we crossed *Gjb2^fl/fl^*mice^11^ to *Tecta-Cre* mice (*Tecta-Cre;Gjb2^fl/fl^,* Cx26 cKO), which exhibit restricted recombination within the sensory epithelium^32^. Immunohistochemical analysis of cochleae from these mice revealed that the large Cx26 gap junction plaque assemblies separating supporting cells in the mature sensory epithelium were eliminated (Fig. 1a and Fig.1b, *white arrows*); however, Cx26 expression was preserved in the outer sulcus, spiral limbus, lateral wall fibrocytes, and stria vascularis (Fig. 1b, *green arrows*), consistent with the restricted expression of *Tecta* (α-Tectorin)^33^. Thus, these conditional Cx26 knockout mice provide a means to explore the specific role of Cx26 within the sensory epithelium.

**Figure 1.**
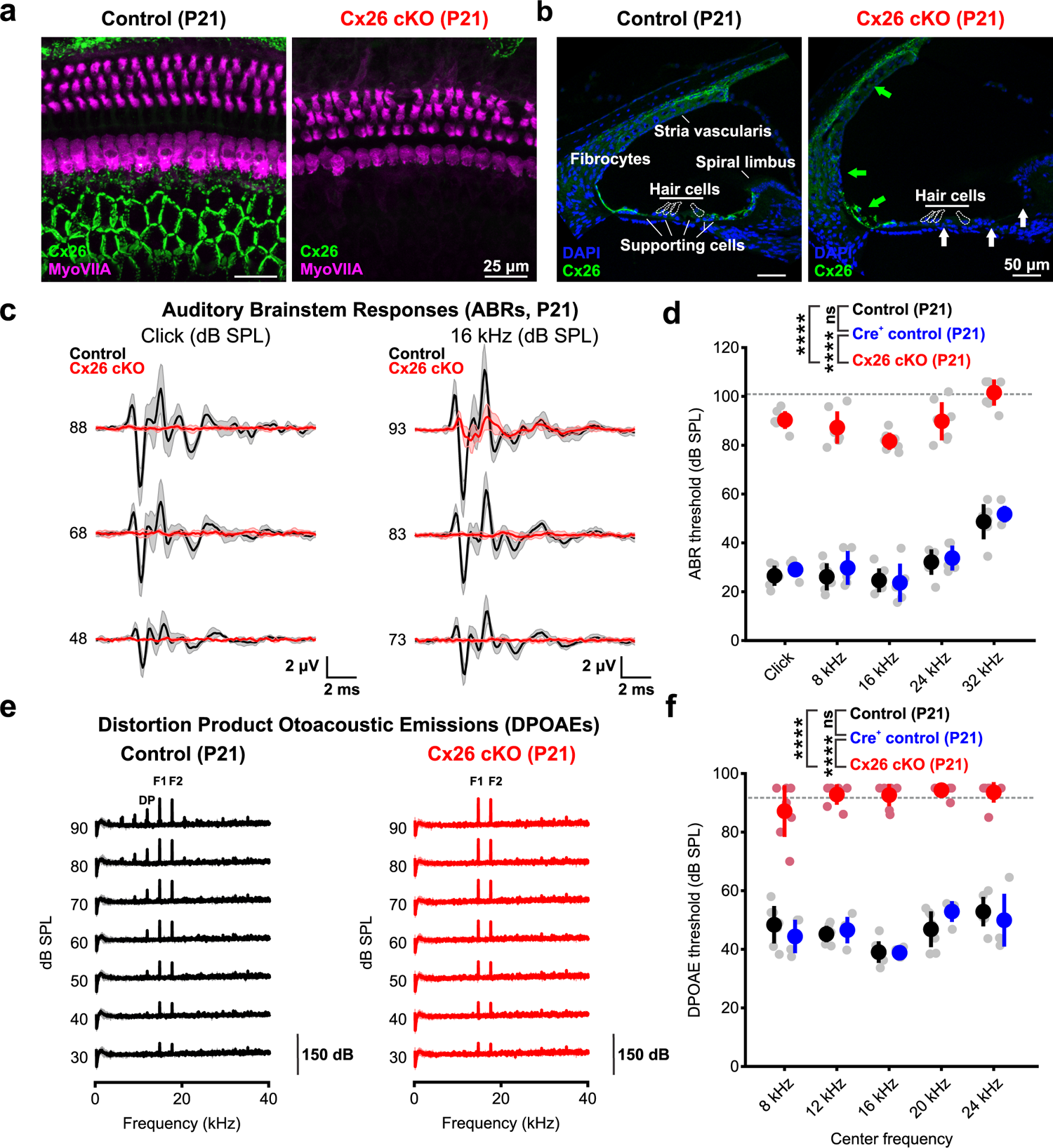
Targeted deletion of Cx26 from the sensory epithelium leads to auditory dysfunction. a) Immunostaining for Connexin 26 (green) in whole mount apical cochlea from P21 control (*Gjb2^fl/fl^*, left) and Cx26 cKO (*Tecta-Cre;Gjb2^fl/fl^*, right) mice. Hair cells (magenta) are labeled by immunoreactivity against MyoVIIA. b) Immunostaining for Connexin 26 (green) in mid-turn cochlea cross section at P21 from control (left) and Cx26 cKO (right) mice. Loss of Cx26 immunostaining is observed in the inner sulcus and supporting cells of the organ of Corti (white arrows) but not within lateral wall fibrocytes or the stria vascularis (green arrows). Hair cell bodies are outlined based on nuclei (DAPI) location. c) Average auditory brainstem response (ABR) traces to broadband click (left) and 16 kHz tone pip (right) stimuli presented at multiple sound pressure levels from control (*Gjb2^fl/fl^*, black, n = 6) and Cx26 cKO (*Tecta-Cre;Gjb2^fl/fl^*, red, n = 7) mice at P21. d) Quantification of P21 ABR thresholds to click and pure tone stimuli in controls (*Gjb2^fl/fl^*, black, n = 6, *Tecta-Cre;Gjb2^fl/+^*, blue, n = 5) and Cx26 cKO (*Tecta-Cre;Gjb2^fl/fl^*, red, n = 7). Grey dashed line indicates maximum speaker output and detection limit. p = 3.4380e-13 (cKO vs control), 1.2801e-9 (cKO vs Cre+ control), 0.4774 (control vs Cre+ control), linear mixed effects model. e) Average recording of distortion product otoacoustic emissions (DPOAEs) for a 16 kHz center frequency at multiple sound pressure levels from control (*Gjb2^fl/fl^*, black, n = 7) and Cx26 cKO (*Tecta-Cre;Gjb2 ^fl/fl^*, red, n = 7) mice at P21. F1; primary tone 1 (14.544 kHz), F2; primary tone 2 (17.440 khz), DP; distortion product. f) Quantification of distortion product thresholds for 5 center frequencies in controls (*Gjb2^fl/fl^*, black, n = 7 and *Tecta-Cre;Gjb2 ^fl/+^*, blue, n = 4) and Cx26 cKO (*Tecta-Cre;Gjb2 ^fl/fl^*, red, n = 7). Grey dashed line indicates maximum speaker output and detection limit. p = 2.3326e-14 (control vs cKO), 4.7233e-10 (Cre+ control vs cKO), 0.9759 (control vs Cre+ control), linear mixed effects model.

To determine if loss of Cx26 from the sensory epithelium is sufficient to disrupt cochlear function and hearing, we recorded auditory brainstem responses (ABRs) and distortion product otoacoustic emissions (DPOAEs) at P21, when normal auditory function has been established. Cx26 cKO mice exhibited profound hearing deficits (Fig. 1c, d), with ABR threshold increases of ∼50-60 dB to both clicks and pure tones relative to controls, in accordance with hearing loss described in mouse models with widespread cochlear Cx26 deletion^12, 34, 35^ and children with loss of function variants in Cx26^13, 14^. Cx26 cKO mice exhibited near-complete loss of DPOAEs at all tested frequencies (Fig. 1e, f), indicating that outer hair cell (OHC) function and cochlear amplification are compromised. However, acoustic sensitivity was not abolished in Cx26 cKO mice, as low amplitude ABRs were still elicited at high sound intensities (> 80 dB SPL), suggesting that mechanotransduction and excitation of SGNs is still possible at this age. These results indicate that removal of Cx26 specifically from supporting cells of the sensory epithelium prior to hearing onset is sufficient to impair later hearing.

Despite this evidence of profound auditory dysfunction, the cochlea in Cx26 cKO mice was largely intact at an age when acoustic sensitivity is normally mature (@P21). The most prominent structural abnormalities in the sensory epithelium were the absence of the Tunnel of Corti and Nuel’s space, resulting in a flattening of the sensory epithelium, as well as subtle morphological changes in hair cell orientation and spacing (Fig. 1a, Extended data Fig. 1), features also observed in mice with broader Cx26 deletion^12, 36^. While no hair cell degeneration was observed prior to ear canal opening (Extended data Fig. 2), partial OHC loss was evident throughout the cochlear length by P21, occurring first in the base (Fig. 1a, Extended data Fig. 2, Extended data Fig. 3a, b, *white arrows*) that became more extensive with age, although this cell loss was lower than reported with cochlea-wide deletion^34, 35^. Remarkably, there was no notable degeneration IHCs and spiral ganglion neurons (SGNs) at P21 in Cx26 cKO mice (Extended data Fig. 3b-d). Progressive degeneration of IHCs eventually occurred with increasing age in a base-apex progression (Extended data Fig. 2). Together, these results demonstrate that selective deletion of Cx26 from supporting cells in the sensory epithelium leads to structural abnormalities in the cochlea and progressive hair cell degeneration following hearing onset.

**Figure 2.**
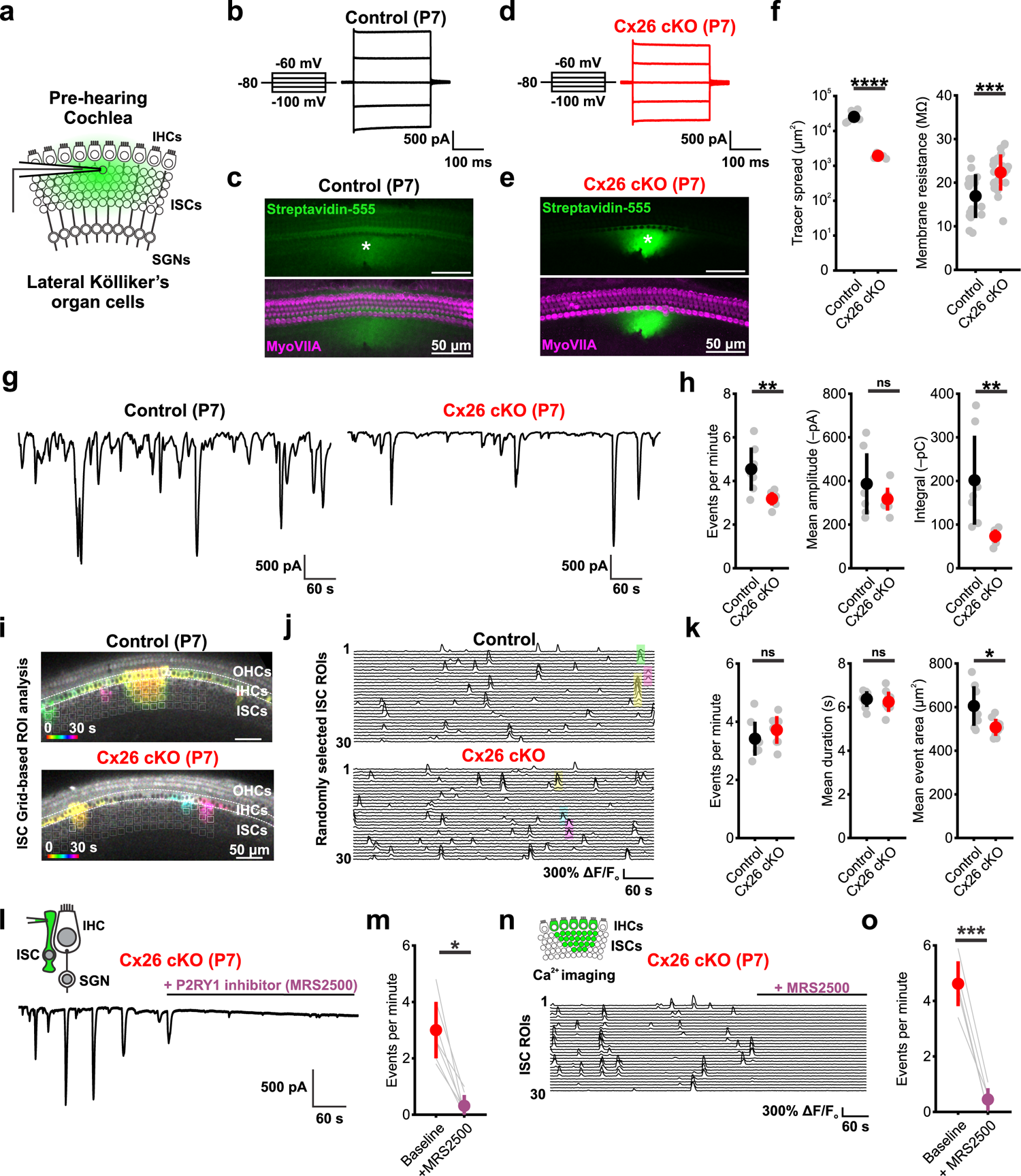
Inner supporting cells generate spontaneous activity in the absence of Cx26. a) Schematic of whole cell patch clamp recording from inner supporting cells (ISCs) with an intercellular tracer within the pipette. b) Current responses elicited by voltage steps within a P7 control (*Gjb2^fl/fl^*) ISC. c) (top) Visualization of intercellular Neurobiotin tracer spread from a P7 control ISC with fluorescent conjugated streptavidin (green). Asterisk indicates patched cell. (bottom) Same as above, but with hair cells labeled by immunoreactivity to Myosin VIIa (magenta). d) Current responses elicited by voltage steps within a P7 Cx26 cKO (*Tecta-Cre;Gjb2^fl/fl^*) ISC. e) Same as (c), but in a Cx26 cKO cochlea. f) Quantification of spatial tracer spread (left) and membrane resistance (right) in ISCs. Tracer spread: n = 4 control, 6 Cx26 cKO cochleae; p = 2.1864e-8, two-sample t-test. Membrane resistance: n = 17 control, 24 Cx26 cKO ISCs; p = 6.3712e-4, two-sample t-test. g) Whole cell voltage clamp recordings of spontaneous activity from control (left) and Cx26 cKO (right) ISCs. h) Quantification of spontaneous event frequency, mean amplitude, and integral (charge transfer). n = 7 control, 8 Cx26 cKO ISCs; p = 0.0045, 0.2405, 0.0061 (frequency, amplitude, integral), two-sample t-test with unequal variances and Benjamini-Hochberg correction for multiple comparisons. i) Temporally pseudocolored 30 s projection of spontaneous calcium transients in ISCs from isolated pre-hearing cochlea from P7 control (*Tecta-Cre;Gjb2^fl/+^;R26-lsl-GCaMP3*, top) and Cx26 cKO (*Tecta-Cre;Gjb2^fl/fl^;R26-lsl-GCaMP3^fl/+^*, bottom) mice. Grid-based region of interest (ROI) analysis is overlaid, with active grids during the 30 s window indicated in white. j) Raster plot of ΔF/F_o_ signals from 30 randomly selected grid ROIs in control (top) and Cx26 cKO (bottom) cochleae. Highlighted transients correspond by color to those indicated in (f). k) Quantification of ISC calcium event frequency (per 0.01 mm^2^), mean duration, and mean event area. n = 8 control, 9 Cx26 cKO; p = 0.3080, 0.5683, 0.0140 (frequency, duration, area), two-sample t-test with Benjamini-Hochberg correction. l) Whole cell voltage clamp recording from a P7 Cx26 cKO ISC with addition of P2RY1 receptor antagonist MRS2500 (1 μM). m) Quantification of spontaneous event frequency at baseline and following P2RY1 antagonism. n = 7 Cx26 cKO ISCs; p = 0.0156, paired Wilcoxon sign rank test. n) Raster plot of calcium ΔF/F_o_ signals from 30 randomly selected ISC grid ROIs from a Cx26 cKO cochlea with addition of MRS2500. o) Quantification of mean event frequency (per 0.01 mm^2^) at baseline and following P2RY1 antagonism. n = 5 Cx26 cKO cochlea; p = 3.0798e-4, paired t-test.

**Figure 3.**
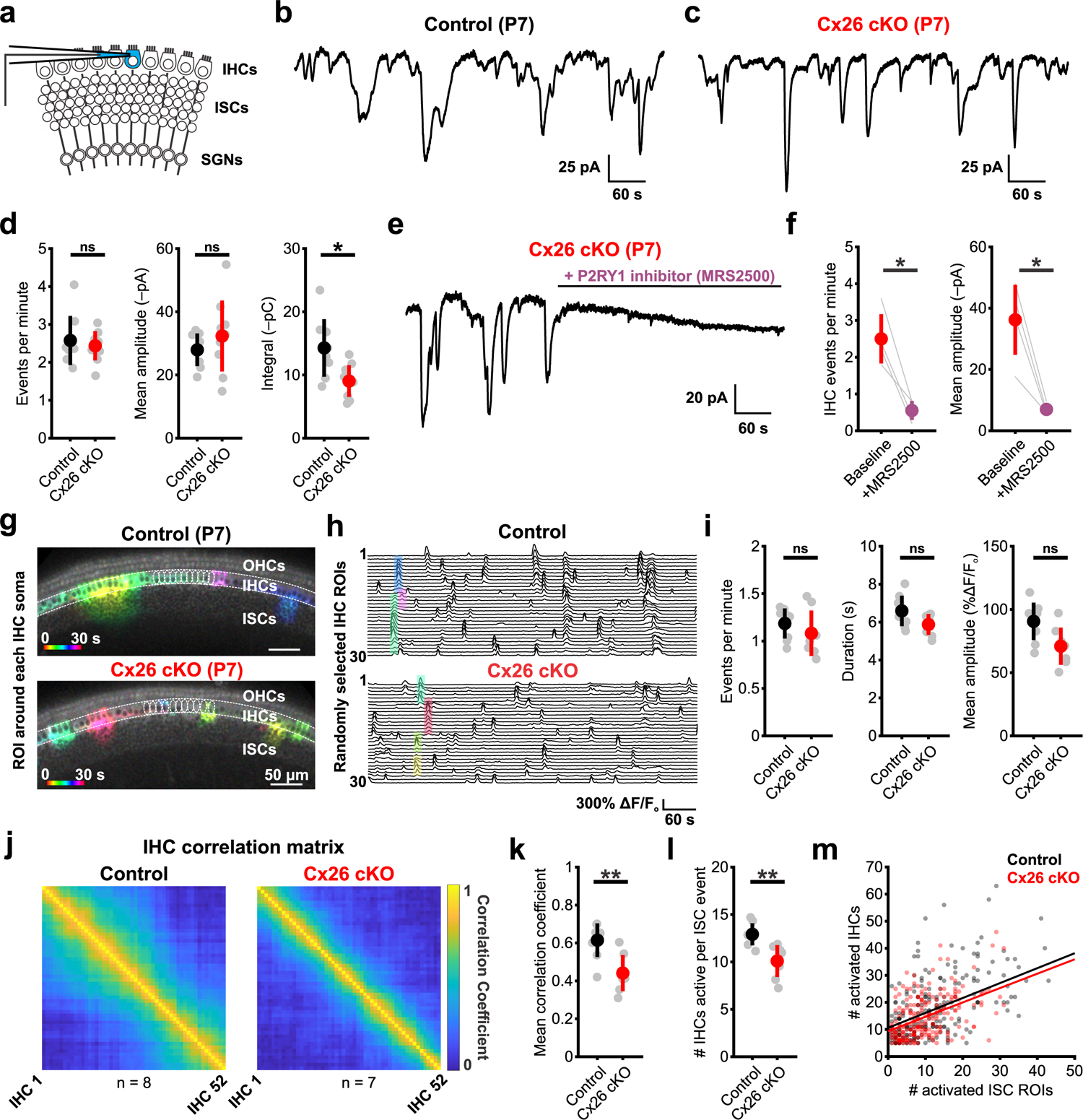
Inner supporting cells coordinate excitation of inner hair cells despite absence of Cx26. a) Schematic of whole cell patch clamp recordings from inner hair cells (IHCs). b) Representative whole cell voltage clamp trace of spontaneous activity from a control (*Gjb2^fl/fl^*) IHC. c) Representative whole cell voltage clamp trace of spontaneous activity from a Cx26 cKO (*Tecta-Cre;Gjb2^fl/fl^*) IHC. d) Quantification of spontaneous inward current frequency, mean amplitude, and integral (charge transfer). n = 8 control, 9 Cx26 cKO IHCs; p = 0.9626, 0.3587, 0.0134 (frequency, amplitude, integral), two-sample t-test with Benjamini-Hochberg correction. e) Whole cell voltage clamp recording from a Cx26 cKO IHC with addition of P2RY1 receptor antagonist MRS2500 (1 μM). f) Quantification of spontaneous inward current frequency and mean amplitude in IHCs before and following addition of MRS2500. n = 4 Cx26 cKO IHCs; p = 0.0132, 0.0168 (frequency, amplitude), paired t-test with Benjamini-Hochberg correction. g) Temporally pseudocolored 30 s projection of spontaneous calcium transients in IHCs in isolated pre-hearing cochlea from control (*Tecta-Cre; Gjb2^fl/+^;R26-lsl-GCaMP3*, top) and Cx26 cKO (*Tecta-Cre;Gjb2^fl/fl^;R26-lsl-GCaMP3*, bottom) mice. Individual IHC oval ROIs are overlaid. h) Raster plot of ΔF/F_o_ signals from 30 randomly selected IHCs in control (top) and Cx26 cKO (bottom) cochleae. Highlighted transients correspond by color to those indicated in (g). i) Quantification of mean IHC calcium event frequency, duration, and amplitude. n = 8 control, 7 Cx26 cKO cochleae; p = 0.4634, 0.1158, 0.0648 (frequency, duration, amplitude). Wilcoxon rank sum test (frequency) or two-sample t-test (duration, amplitude) with Benjamini-Hochberg correction. j) Mean correlation matrix of IHC ΔF/F_o_ signals in control (left) and Cx26 cKO (right) cochleae. k) Quantification of mean correlation coefficient (80^th^ percentile) between IHCs. n = 7 control, 7 Cx26 cKO cochleae; p = 0.0048, two-sample t-test. l) Quantification of mean number of IHCs active per supporting cell calcium event. n = 7 control, 7 Cx26 cKO cochleae; p = 0.0057, two-sample t-test. m) Scatter plot of activated inner supporting cell (ISC) grid ROIs and activated IHCs during supporting cell calcium events in control (grey) and Cx26 cKO (pink) cochleae. Solid line indicates linear regression model of group data in control (black) and Cx26 cKO (red).

### ISCs continue to generate spontaneous activity in the absence of Cx26

Supporting cells in the developing cochlea express several distinct gap junction proteins, most prominently Cx26 and Cx30, which can form intercellular channels and uncoupled hemichannels at the cell surface. Notably, expression of these two gap junctions is closely linked and disruption of Cx26 has been shown to disrupt Cx30 expression, and vice versa^34, 37, 38^. To determine how loss of Cx26 affects the physiological properties and intercellular coupling of ISCs, we recorded from ISCs in cochleae acutely isolated at P7, targeting cells in the lateral region of Kölliker’s organ that are responsible for generating spontaneous activity^18, 19^ (Fig. 2a). ISCs in control cochleae exhibited both low membrane resistance (16.9 ± 5.2 MΩ) (Fig. 2b) and extensive coupling, visible as the diffuse spread of tracer throughout Kölliker’s organ^3^ (Fig. 2c, *green fluorescence*). Unexpectedly, ISCs in Cx26 cKO mice also exhibited extensive tracer spread (Fig. 2d, e) and retained low membrane resistance (22.3 ± 4.1 MΩ) (Fig. 2f), indicating that they remain coupled through gap junctions; however, tracer movement was more restricted, with less diffusion within Kölliker’s organ and between inner phalangeal cells and inner pillar cells. Thus, even in the absence of Cx26, ISCs in the developing cochlear epithelium remain coupled through gap junctions.

ISCs exhibit spontaneous electrical activity from the late embryonic period through to ear canal opening, after which it rapidly disappears. Spontaneous activity is initiated by periodic release of ATP and activation of P2RY1 autoreceptors on ISCs^39, 40^. To explore how the absence of Cx26 influences ISC activity prior to hearing onset, we recorded spontaneous activity patterns from ISCs using whole cell recording and imaged intracellular calcium changes from the cochlear sensory epithelium in acutely excised cochleae from P7 mice lacking Cx26 (Extended data Fig. 4a, b). Periodic inward currents were still visible in ISCs from Cx26 cKO mice, but they were less frequent and shorter in duration than in controls, leading to less overall charge transfer (integral of the current trace) (Fig. 2g, h). These reductions are consistent with impaired electrical coupling, as activity throughout the supporting cell syncytium is detected when recording from individual cells^3^. Calcium imaging in the cochlea (*Tecta-Cre;Rosa26-lsl-GCaMP3 ± Gjb2^fl/fl^*)^41^ revealed that spontaneous ISC calcium transients from P7 Cx26 cKO cochlea occurred at a similar frequency as controls, and had a comparable average duration, but extended over a slightly smaller area along both medial-lateral and tonotopic axes (Fig. 2i-k, Supplementary Video 1). Both inward currents and calcium transients in Cx26 cKO cochleae were blocked by MRS2500 (1µM)^32^, a P2RY1 selective antagonist (Fig. 2l-o), indicating that ISCs release ATP and initiate a P2RY1-mediated signaling cascade in the absence of Cx26. Moreover, osmotic shrinkage (crenation) of ISCs, which follows ATP-dependent ion efflux^19^, was also evident in Cx26 cKO ISCs (Extended data Fig. 4c). The average spread of these osmotic crenations was reduced in Cx26 cKO cochleae, consistent with the reduced extent of ISC calcium transients along the tonotopic axis (Fig. 2k, Extended data Fig. 4d). Together, these data indicate that key features of supporting cell spontaneous activity in the developing cochlea (ATP release, coordinated calcium transients, and ionic flux from ISCs) are preserved when Cx26 is absent from the supporting cell gap junction network.

**Figure 4.**
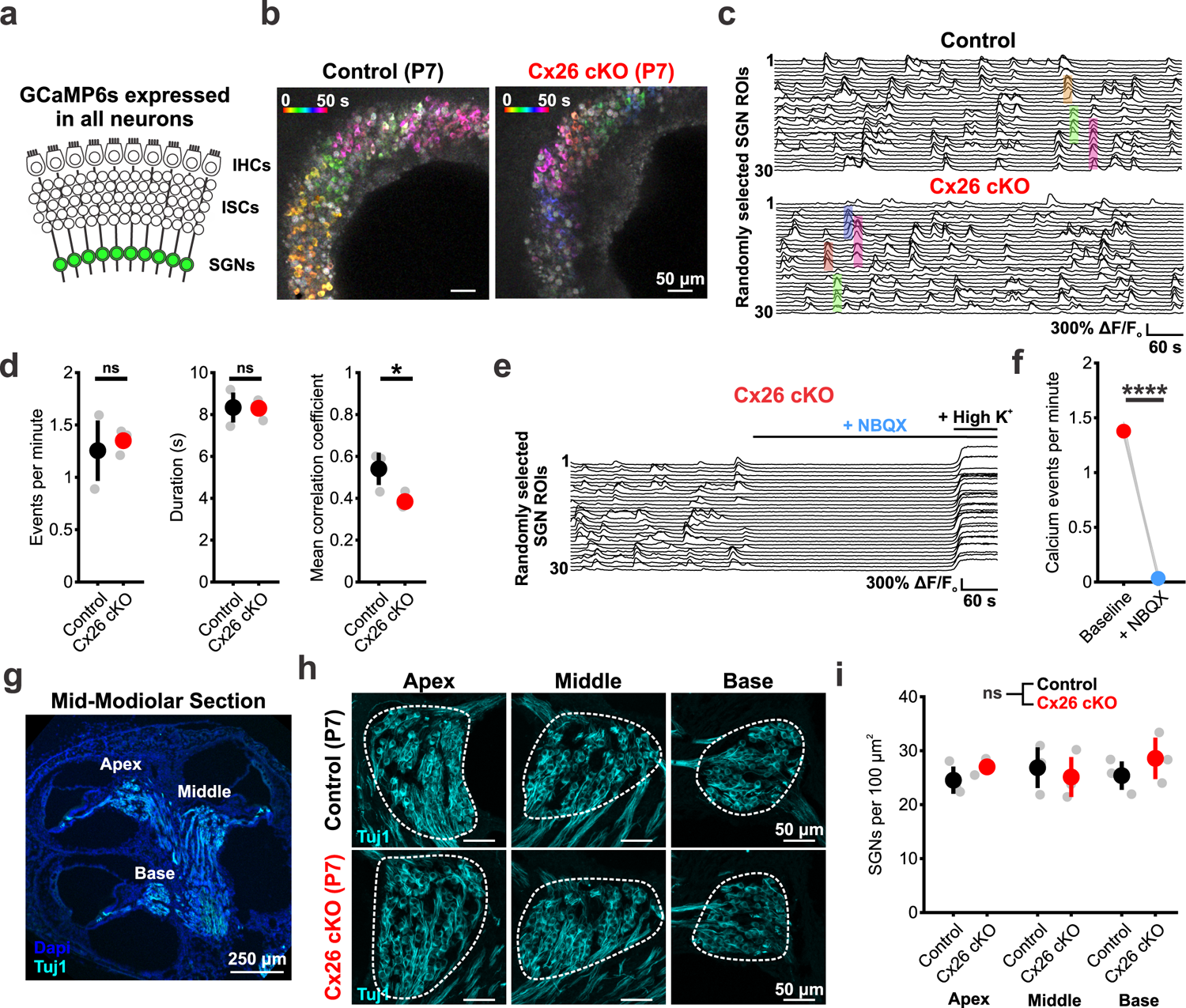
Spontaneous synaptic activation of spiral ganglion neurons persists in the absence of supporting cell Cx26. a) Schematic of calcium imaging in spiral ganglion neurons (SGNs) in *Snap25-T2A-GCaMP6s* mice. b) Temporally pseudocolored 50 s projection of coordinated spontaneous calcium transients in SGNs from isolated pre-hearing cochlea in control (*Gjb2^fl/fl^;Snap25-T2A-GCaMP6s*, top) and Cx26 cKO (*Tecta-Cre;Gjb2^fl/fl^;Snap25-T2A-GCaMP6s*, bottom). c) Raster plot of ΔF/F_o_ signals from 30 randomly selected SGN ROIs in control (top) and Cx26 cKO (bottom) cochleae. Highlighted transients correspond by color to those indicated in b. d) Quantification of spontaneous SGN calcium transient frequency, mean amplitude, and mean duration. n = 3 control, 5 Cx26 cKO cochleae; p = 0.7857, 0.9345, 0.0114 (frequency, duration, correlation), two-sample t-test with Benjamini-Hochberg correction. e) Raster plot of ΔF/F_o_ signals from 30 randomly selected SGN ROIs in a Cx26 cKO cochlea with addition of NBQX (50 μM). High potassium (High K^+^) artificial cerebrospinal fluid (aCSF) is added following NBQX to ensure intact calcium responses. e) Quantification of calcium event frequency within Cx26 cKO SGNs with addition of NBQX. n = 4 Cx26 cKO cochleae; p = 1.8187e-5, paired t-test. f) (left) Low magnification image of spiral ganglion neurons (SGNs) labeled by immunoreactivity to Tuj1 (cyan) in mid-modiolar cross section of P7 cochlea. Labels indicate locations of apical, middle, and basal SGN counts. (right) Representative high-magnification images of SGN soma in apical, middle, and basal cochlea from control (*Gjb2^fl/fl^*, top) and Cx26 cKO (*Tecta-Cre;Gjb2^fl/fl^*, bottom) at P7. Dashed lines indicate SGN compartment used for area measurement. g) Quantification of SGN density in apical, middle, and basal cochlea at P7. n = 3 control, 3 Cx26 cKO; p = 0.5885, linear mixed effects model.

### ISCs coordinate excitation of IHCs and SGNs in the absence of Cx26

In the developing cochlea, ISC mediated extrusion of potassium leads to depolarization of nearby IHCs within a restricted region of the organ of Corti^19, 40^. To determine whether ISCs in Cx26 cKO mice remain capable of eliciting IHC depolarization, we monitored the spontaneous activity of IHCs in acutely isolated P7 cochleae (Fig. 3a). Consistent with persistence of large ATP-induced inward currents in ISCs, IHCs in Cx26 cKO mice exhibited spontaneous inward currents at a frequency and amplitude comparable to controls (Fig. 3b-d). In accordance with the briefer ISC events, IHC events were also shorter in duration, leading to a decrease in total charge transfer (integral) per event (Fig. 3d). This spontaneous activity was similarly dependent on ISC ATP release, as it was abolished by MRS2500 (Fig. 3e, f). Notably, the physiological properties of IHCs (current-voltage relationship, resting membrane potential, and input resistance) were comparable between control and Cx26 cKO mice (Extended data Fig. 5), suggesting that these subtle changes IHC spontaneous activity are due to extrinsic changes in supporting cell ion flux surrounding IHCs.

**Figure 5.**
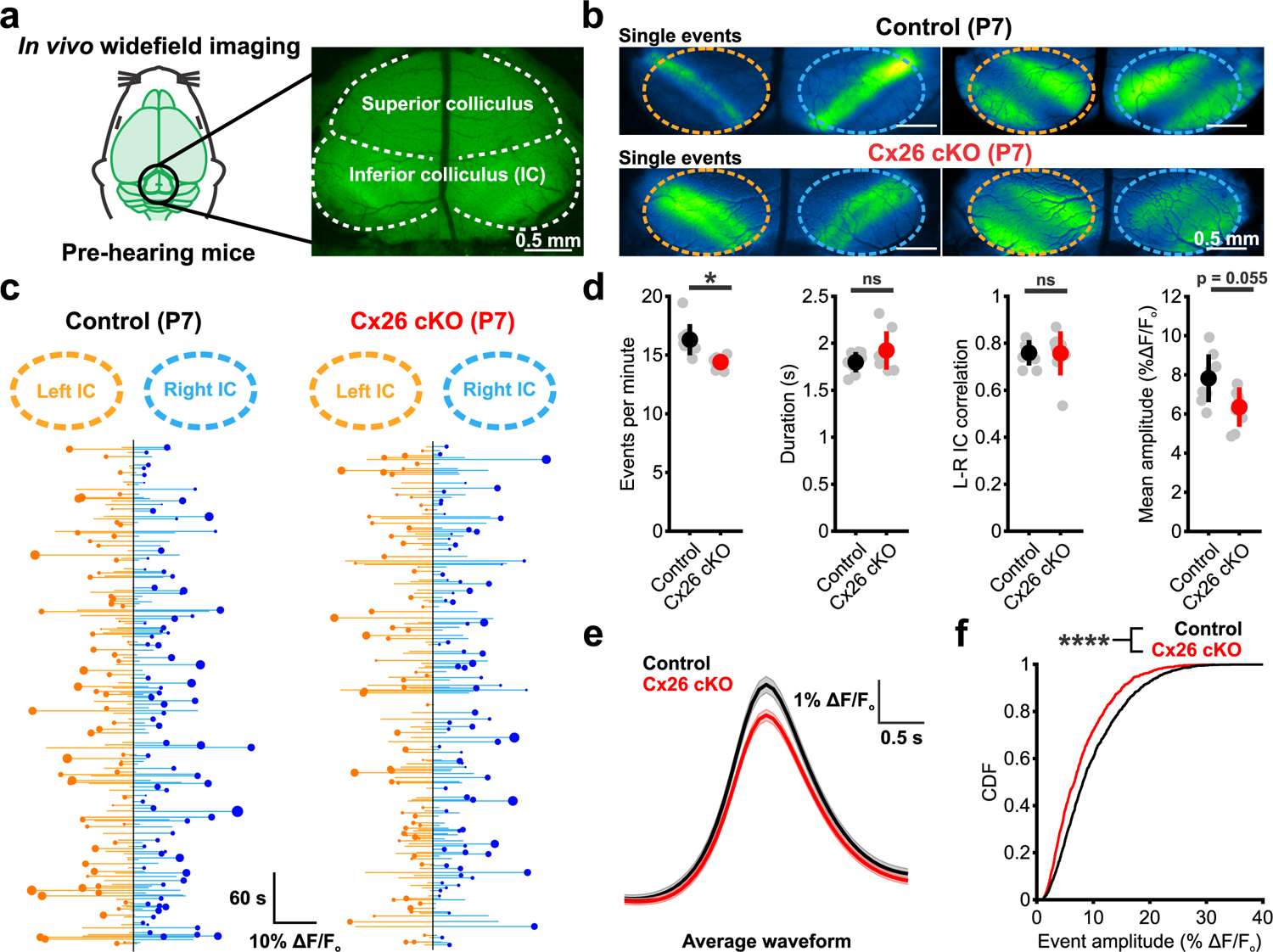
Spontaneous activity persists in the developing auditory system of Cx26 cKO mice. a) Schematic of *in vivo* widefield epi-fluorescent imaging paradigm to visualize pre-hearing neural activity in the inferior colliculus (IC). b) Representative single calcium events in P7 inferior colliculus from control (*Gjb2^fl/fl^;Snap25-T2A-GCaMP6s*, top) and Cx26 cKO (*Tecta-Cre;Gjb2^fl/fl^;Snap25-T2A-GCaMP6s*, bottom) mice occurring within isofrequency bands along the future tonotopic axis of the IC. c) Trace of spontaneous calcium transients over 10 minutes in the right and left inferior colliculi from control (left) and Cx26 cKO (right) mice. Each line represents a calcium event, circles indicate which colliculus (right vs. left) exhibited a larger amplitude during bilateral events, with the size of the circle indicating the relative difference in amplitude between right and left, with larger circles representing more asymmetric bilateral events. d) Quantification of spontaneous calcium event frequency, duration, degree of correlation between right and left colliculi, and mean event amplitude. n = 8 control, 8 Cx26 cKO mice; p = 0.0122, 0.2287, 0.9607, 0.0548 (frequency, duration, correlation, amplitude), two-sample t-test with Benjamini-Hochberg correction. e) Waveform of the mean spontaneous event aligned to peak amplitude from control and Cx26 cKO mice. n = 1315 events from 8 control mice, 1161 events from 8 Cx26 cKO mice. f) Cumulative distribution plot of peak event amplitude. n = 1315 events from 8 control mice, 1161 events from 8 Cx26 cKO mice; p = 2.8281e-11, two-sample Kolmogorov-Smirnov test.

As fewer ISCs are activated by each ATP release event in Cx26 cKO cochleae (Fig. 2k) and currents induced in IHCs had less charge transfer, we hypothesized that this may result in more spatially restricted activation of IHCs. Calcium imaging revealed that transients within individual IHCs occurred at a similar rate in control and cKO cochlea, and the duration and amplitude of these events were not statistically different, although they trended lower (Duration: p = 0.12; Amplitude: p = 0.065; two-sample t-test with Benjamini-Hochberg FDR correction) (Fig. 3h, i). IHC activity correlation matrices revealed that the extent of synchrony was lower in Cx26 cKO cochleae (Fig. 3j, k), with ∼20% fewer IHCs activated per ISC calcium transient (Fig. 3l). This reduction in IHC activity in Cx26 cKO mice was highly correlated with the extent of ISC activation, as linear regression fits to the number of IHCs and ISCs activated per event had comparable slopes (Fig. 3m), suggesting that the reduction in spread of activity among ISCs in Cx26 cKO mice, and thus the area from which potassium is released, leads to fewer IHCs being depolarized.

IHCs in the developing cochlea are excitable and can generate calcium action potentials upon depolarization, resulting in glutamate release onto SGN afferent fibers, activation of AMPA and NMDA receptors, depolarization, and burst firing^21, 42^. To determine if synaptic connectivity between IHCs and SGNs is preserved in the absence of Cx26, we monitored SGN activity using calcium imaging in excised P7 cochlea from control (*Gjb2^fl/fl^*;*Snap25-T2A-GCaMP6s*) and Cx26 cKO mice (*Tecta-Cre;Gjb2^fl/fl^*;*Snap25-T2A-GCaMP6s*)^43^ (Fig. 4a). In cochleae from Cx26 cKO mice, spontaneous calcium transients occurred periodically among groups of SGNs located within spatially restricted regions of the cochlea at similar rates and duration as controls, but with reduced correlation (Fig. 4b-d). SGN calcium transients in Cx26 cKO cochleae were similarly dependent on synaptic excitation by IHCs, as they were blocked by the AMPA-receptor antagonist NBQX (50 μM) (Fig. 4e, f, Supplementary Video 2). Electrical activity of SGNs during this critical period of development has been shown to influence both the maturation and survival of SGNs^44, 45^; consistent with preservation of activity, Tuj1 immunoreactivity revealed that SGNs in P7 Cx26 cKO mice were present in numbers comparable to control mice (Fig. 4g-i). Together, these results indicate that there is unexpected preservation of IHC-SGN synaptic coupling and spontaneous activity patterns in the cochleae of Cx26 cKO mice prior to hearing onset, resulting in coordinated activation of neurons that project to central auditory centers.

### Spontaneous activity persists in the developing auditory system of Cx26 cKO mice

Action potential bursts generated in SGNs propagate along the ascending auditory pathway with remarkable fidelity, inducing correlated firing of neurons within isofrequency zones in central auditory centers^21, 22, 46^. In the inferior colliculus (IC), this peripheral input manifests as discrete bands of neuronal activity aligned to the future tonotopic axis^22, 47^. To determine if the correlated firing of auditory neurons within these lamina is affected in Cx26 cKO mice, we performed widefield neuronal calcium imaging in awake, unanesthetized mice (*Gjb2^fl/fl^*;*Snap25-T2A-GCaMP6s* and *Tecta-Cre;Gjb2^fl/fl^*;*Snap25-T2A-GCaMP6s*) through a cranial window placed over the midbrain (Fig. 5a). Consistent with persistence of SGN burst firing in P7 Cx26 cKO mice, temporally and spatially correlated bands of neuronal activity were observed within discrete domains of the IC, resembling patterns observed in controls (Fig. 5b, Supplementary Video 3, Extended data Fig. 6a-c). Plots of event amplitude across both lobes of the IC revealed that spontaneous activity in both mouse lines was highly variable in both amplitude (Fig. 5c, length of *horizontal orange and blue lines*) and left-right hemisphere dominance (Fig. 5c, size of *orange and blue circles*), mirroring the stochastic nature of ATP release events in the cochlea. Overall, events were slightly less frequent and trended lower in mean amplitude in Cx26 cKO mice (Fig. 5d); however, the bilateral nature and temporal kinetics (half-width) of these events were unchanged. Consistent with the trend towards smaller mean event amplitude, the average response peak was smaller and the cumulative distribution of event amplitudes and spatial integral were slightly shifted towards smaller events in Cx26 cKO mice, suggesting that fewer neurons were activated synchronously during each event (Fig. 5e, f, Extended data Fig. 6d). While differences in the width of spontaneous events along the future tonotopic axis were not observed (Extended data Fig. 6e), analysis of spontaneous events at single-cell resolution revealed subtle decreases in spatial activation of neurons in Cx26 cKO mice (Extended data Fig. 6f, g), consistent with observations in the isolated cochlea. Together, these *in vivo* results indicate that highly correlated bursts of neural activity continue to be induced in central auditory neurons prior to hearing, despite the lack of Cx26 expression within the organ of Corti.

**Figure 6.**
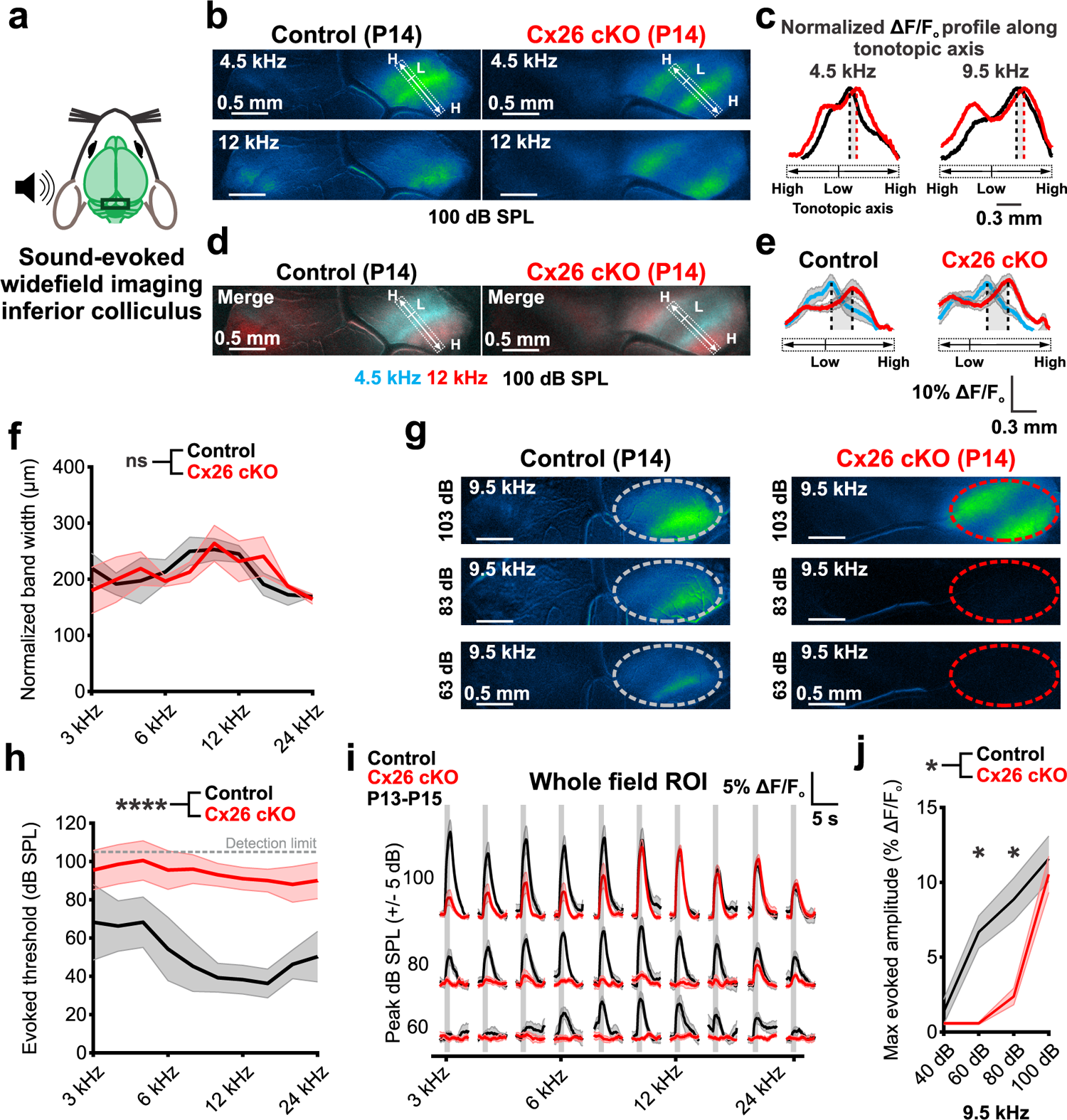
Reduced acoustic sensitivity but retained midbrain tonotopic organization in Cx26 cKO mice. a) *In vivo* widefield imaging of tone-evoked inferior colliculus (IC) neural activity in unanesthetized mice after hearing onset. b) Tone-evoked neural calcium transients in IC from P14 control (*Gjb2^fl/fl^;Snap25-T2A-GCaMP6s*, left) and P14 Cx26 cKO (*Tecta-Cre;Gjb2^fl/fl^;Snap25-T2A-GCaMP6s*, right) mice at 100 dB sound pressure level (SPL). Rectangular ROIs were placed along the tonotopic axis of the contralateral IC (low (L) to high (H) frequency), perpendicular to pure tone evoked bands, to determine the peak response location for a pure tone. c) Plot of tone-evoked normalized mean fluorescence along the tonotopic axis of the IC in control and Cx26 cKO mice. Dashed lines indicate location of peak pure tone response along tonotopic axis, gray shading indicates shift in peak response location for a given pure tone between control and Cx26 cKO mice. n = 4 control mice, 5 Cx26 cKO mice. d) Pseudocolored tone-evoked calcium transients depicting spatial segregation of low and higher frequency stimuli along tonotopic axis. e) Plot of tone-evoked mean fluorescence along the tonotopic axis of the IC in control (left) and Cx26 cKO (right) mice. Dashed lines indicate location of peak response along tonotopic axis, gray shading indicates spatial separation in peak response location for 4.5 (cyan) and 12 kHz (red) pure tones in control and Cx26 cKO mice. f) Quantification of pure tone evoked spatial activation (band width, 75^th^ percentile) normalized to peak fluorescence response amplitude along the tonotopic axis of the IC at 100 dB SPL. n = 4 control, 5 Cx26 cKO mice; p = 0.7575, linear mixed effects model. g) IC neural calcium transients to a 9.5 kHz stimulus from 103 to 63 dB sound pressure level (SPL) in a control and Cx26 cKO mouse. Circle in right IC depicts ROI for subsequent quantification of threshold and amplitude. h) Quantification of pure tone sound-evoked thresholds. n = 4 control, 10 Cx26 cKO mice; p = 1.6305e-6, linear mixed effects model. i) Quantification of tone-evoked fluorescence in IC across a range of frequency and sound level stimuli. Vertical gray bar indicates tone presentation. n = 6-7 control mice, 9-10 *Tmem16a* cKO mice, mean ± SEM. j) Rate-level functions characterizing maximum whole IC response amplitude at 9.5 kHz. mean ± SEM, n = 6-7 control mice, 9-10 *Tmem16a* cKO mice; p = 0.0166, linear mixed effects model with Sidák post hoc test.

### Cx26 cKO mice exhibit reduced acoustic sensitivity but normal topographic organization

Despite the preservation of synaptic connections between IHCs and SGNs and robust spontaneous activity during early development in Cx26 cKO mice, these mice exhibited extreme auditory dysfunction at P21 (Fig. 1). To determine if this functional deficit is present soon after hearing onset or develops later, we imaged sound-evoked neural activity within central auditory nuclei in unanesthetized mice just after ear canal opening (P13-P15) (Fig. 6a), an approach that is more sensitive than ABR, as detection does not require synchronous activation of large numbers of neurons and can be recorded without anesthesia. In control mice, unilateral sound stimuli (3-24 kHz sinusoidally amplitude-modulated pure tones) at a range of intensities (40-100 dB SPL) elicited neuronal activation within isofrequency domains of the IC (Fig. 6b)^22^. Unexpectedly, stimulation with pure tones at > 90 dB SPL also induced robust, tonotopically-restricted activation of neurons within the IC of Cx26 cKO mice, allowing assessment of the functional organization of these regions. Plotting evoked fluorescence profile along the tonotopic axis of the IC revealed that while the best frequency locations to pure tone stimuli were slightly shifted along the tonotopic axis towards higher frequency (lateral) regions in Cx26 cKO mice (Fig. 6c, Extended data Fig. 7a, b), proper spatial separation and normal width of isofrequency lamina were maintained (Fig. 6d-f, Extended data Fig. 7c). In accordance with ABR recordings at this age (Extended data Fig. 8a, b), pure tone response thresholds were markedly elevated in Cx26 cKO mice relative to controls (Fig. 6g, h); however, at suprathreshold sound intensities, the average amplitudes and time courses of evoked calcium responses were comparable those recorded from control mice at 8 kHz and higher (Fig. 6i-j, Extended data Fig. 6d, e). Notably, these elevations in auditory thresholds preceded substantial hair cell degeneration (Extended data Fig. 8c, d), raising the possibility that structural alterations in the organ of Corti prevent proper amplification of sound induced vibrations (Extended data Fig. 7e, f).

**Figure 7.**
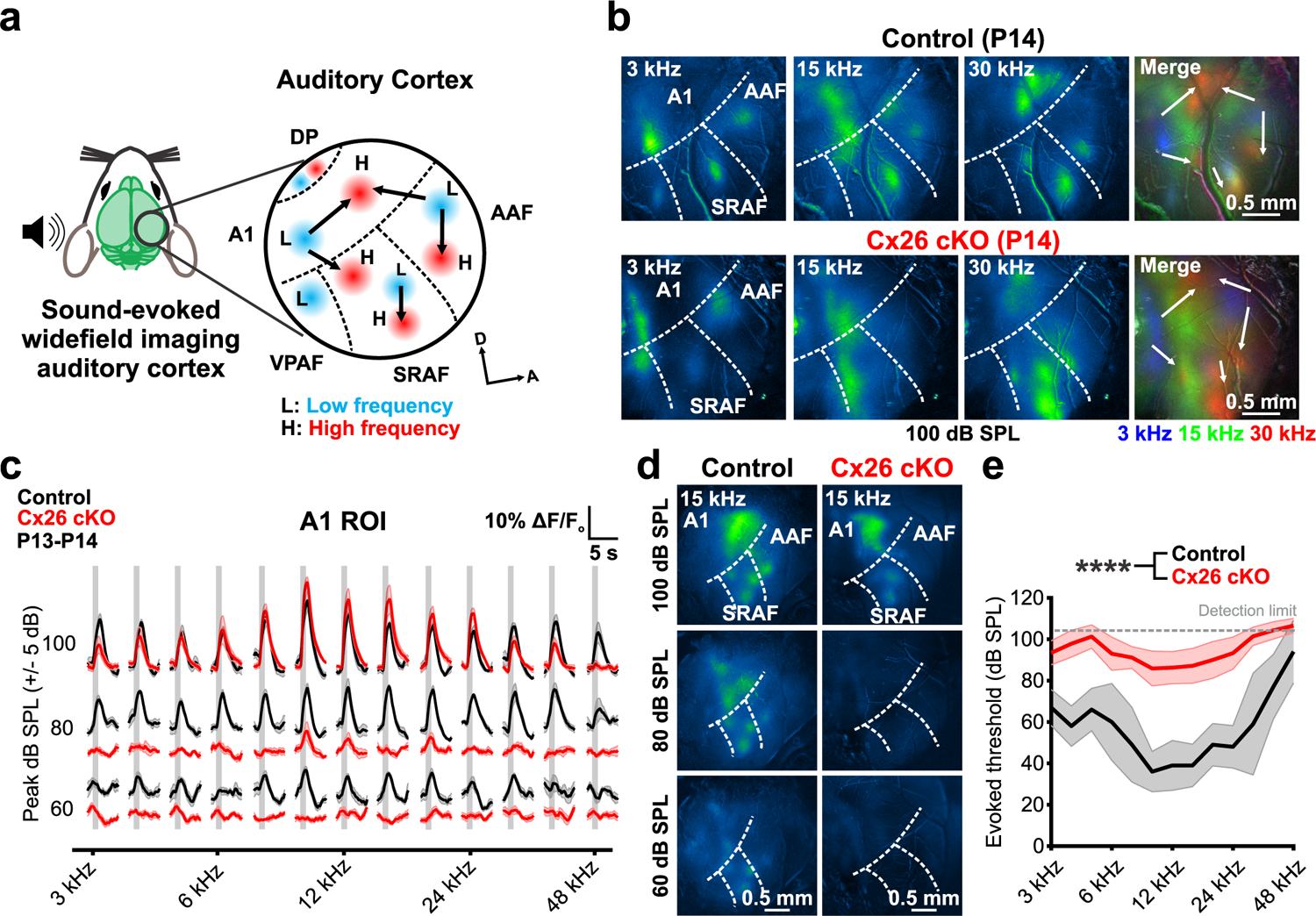
Suprathreshold stimuli elicit tonotopically organized calcium responses in auditory cortex of immature Cx26 cKO mice. a) Schematic depicting tonotopic organization of mouse auditory cortex, adapted from ^48, 50^. L: low frequency, H: high frequency, A1: primary auditory cortex, AAF: anterior auditory field, SRAF: suprarhinal auditory field, VPAF: ventral posterior auditory field, DP: dorsal posterior. b) Tone-evoked widefield neural calcium transients in AC from P14 control (*Gjb2^fl/fl^;Snap25-T2A-GCaMP6s*) and P14 Cx26 cKO (*Tecta-Cre;Gjb2^fl/fl^;Snap25-T2A-GCaMP6s*) mice at 100 dB SPL. Merged image shows tonotopic segregation of pseudocolored pure tone responses from low (L) to high (H) frequency along tonotopic axes. c) Quantification of tone-evoked fluorescence in A1 across a range of frequency and sound level stimuli in P13-P14 control and Cx26 cKO mice. Vertical gray bar indicates tone presentation. n = 5 control mice, 7 *Tmem16a* cKO mice, mean ± SEM. d) AC neural calcium transients to a 15 kHz pure tone stimulus from 100 to 60 dB SPL in a control and Cx26 cKO mouse. e) Quantification of pure tone sound-evoked thresholds. n = 5 control, 7 Cx26 cKO mice; p = 8.8449e-6, linear mixed effects model.

**Figure 8.**
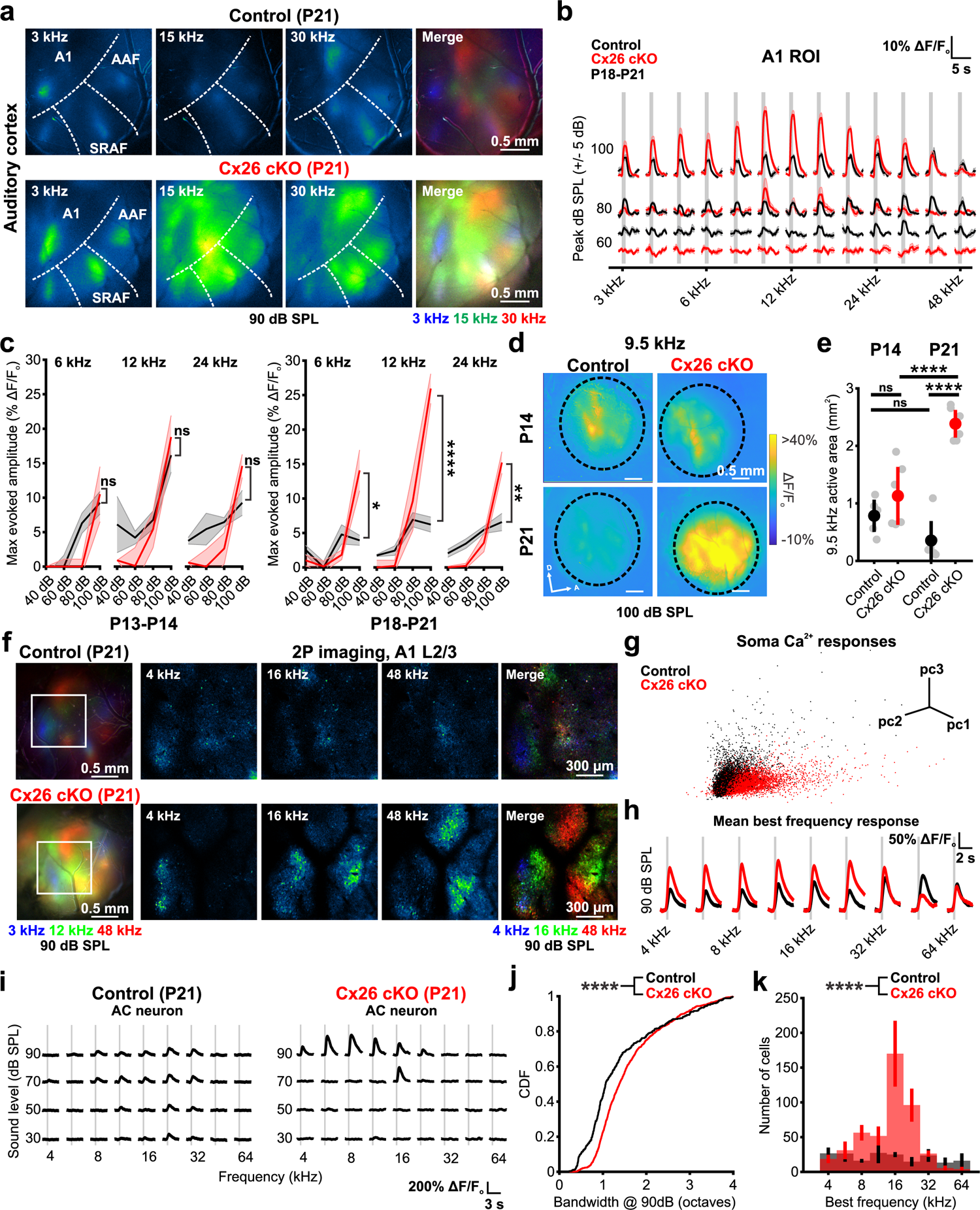
Rapid increases in central gain after hearing onset in Cx26 cKO mice. a) Tone-evoked widefield neural calcium transients in AC from P21 control (*Gjb2^fl/fl^;Snap25-T2A-GCaMP6s*) and P21 Cx26 cKO (*Tecta-Cre;Gjb2^fl/fl^;Snap25-T2A-GCaMP6s*) mice at 90 dB SPL. Merged image shows tonotopic segregation of pseudocolored pure tone responses along tonotopic axes. b) Quantification of tone-evoked fluorescence in A1 across a range of frequency and sound level stimuli in P18-P21 control and Cx26 cKO mice. Vertical gray bar indicates tone presentation. n = 7 control mice, 9 *Tmem16a* cKO mice, mean ± SEM. c) Rate-level functions characterizing maximum A1 response amplitude at 6, 12, and 24 kHz at P13-P14 (left) and P18-P21 (right). Mean ± SEM, n = 7 P18-P21, 5 P13-P14 control mice, 9 P18-P21, 7 P13-P14 Cx26 cKO mice; P13-P14: p = 0.7551, 0.5816, 0.2418, Wilcoxon rank sum test (6 kHz) or two-sample t-test (12 kHz, 24 kHz) with Benjamini-Hochberg correction. P18-P21: p = 0.0164, 2.1097e-7, 0.0026, Wilcoxon rank sum test (6 kHz) or two-sample t-test (12 kHz, 24 kHz) with Benjamini-Hochberg correction. d) Normalized fluorescence responses in auditory cortex (circled) of control and Cx26 cKO mice at P14 and P21 to a 9.5 kHz pure tone stimulus at 100 dB SPL. e) Quantification of activated area in auditory cortex in response to a 9.5 kHz pure tone. n = 5 P13-P14, 6 P16-P21 control mice, 6 P13-P14, 7 P16-P21 Cx26 cKO mice; p = 5.8256e-8, one way ANOVA with Tukey’s HSD post hoc comparisons. f) (left) Widefield macroscopic tonotopic maps of auditory cortex in a P21 control (top) and Cx26 cKO (bottom). White square indicates A1 region for two photon imaging. (right) Sound-evoked neuronal fluorescence responses to pure tones at 90 dB SPL in layer II/III of A1 from mouse at left. g) Principal component analysis based on the maximum fluorescence amplitudes of all neural soma in A1 following tonal stimuli across all presented frequency and intensity levels. Each dot represents tone-evoked calcium responses of a neural soma. h) Mean fluorescence changes of all tone-responsive neurons sorted by their best frequency stimulus at 90 dB SPL. i) Fluorescence changes in a representative A1 neuron to a range of frequency and intensity stimuli in a control and Cx26 cKO mouse. j) Cumulative distribution of gaussian bandwidth at 90 dB SPL. n = 432 control cells (3 mice), 1088 Cx26 cKO cells (3 mice); p = 1.8327e-14, two-sample Kolmogorov-Smirnov test. k) Histogram of mean number of tone-responsive neurons binned by their best frequency. n = 567 control cells (3 mice), 1379 Cx26 cKO cells (3 mice); p = 7.3571e-15, two-sample Kolmogorov-Smirnov test.

To assess whether connections between the midbrain and thalamus enable transmission of sound evoked activity to the nascent auditory cortex (AC), we installed acute cranial windows over the AC and imaged sound-evoked responses without anesthesia just after ear canal opening (P13-P14, Fig. 7a). In response to tones of different stimulus intensities, control mice exhibited several discrete foci of neuronal activity within primary (A1), suprarhinal (SRAF), and anterior (AAF) auditory cortical regions^48–50^ (Fig. 7b), consistent with the early establishment of tonotopic partitioning of acoustic information^51, 52^. In response to loud tones, Cx26 cKO mice also exhibited sound-evoked neuronal activation with appropriate tonotopic organization that were comparable in amplitude to controls at similar sound intensities (Fig. 7b, c); however, in accordance with decreased sensitivity observed in the IC, thresholds for neuronal responses in AC at these early ages were also much higher in Cx26 cKO mice (Fig. 7d, e). Together, these results indicate that mice deprived of Cx26 in the organ of Corti retain proper tonotopic organization within auditory centers following ear canal opening, but exhibit a profound deficit in acoustic sensitivity

### Rapid increases in central gain after hearing onset in Cx26 cKO mice

Neural circuits within mouse central auditory centers adapt to the acoustic environment within a critical period in the third postnatal week^53–55^. We leveraged the increased sensitivity of *in vivo* neuronal calcium imaging to assess how deprivation of normal acoustic input during this key developmental phase alters acoustic response characteristics. Similar to the responses in mice recorded just after hearing onset, pure tones at high intensity stimuli (80-100 dB SPL) elicited neuronal activation within auditory cortex of P21 Cx26 cKO mice (Fig 8a); however, while response amplitudes decreased between P14 and P21 in control mice, consistent with emerging local inhibition^51, 56, 57^, Cx26 cKO mice exhibited an opposite trend, with evoked response amplitudes to suprathreshold stimuli increasing dramatically during this period (Fig. 8a, b), resulting in steeper intensity-level functions (Fig. 8b, c). This increased fluorescence was accompanied by broader spatial activation to pure tone responses, particularly within the mid-frequency range (Fig. 8d, e). *In vivo* two photon imaging from Cx26 cKO mice revealed that individual cortical neurons within AC exhibited distinct pure tone activation patterns from control cells, with larger calcium transients to a given best frequency suprathreshold stimulus (Fig. 8f-h). Broader spatial activation could result from a wider frequency sensitivity of AC neurons; however, we observed only a modest increase in the bandwidth of sharply tuned AC neurons in Cx26 cKO mice (Fig. 8i, j). Rather, there was a dramatic increase in the number of AC neurons engaged by suprathreshold pure tones (9 frequencies, 4-64 kHz, 30-90 dB SPL) in Cx26 cKO mice (Activated neurons: control, 6.9 ± 2.7%; Cx26 cKO, 13.0 ± 4.4%) across a wider range of cortical territory, particularly for neurons tuned to mid-frequency stimuli (Fig. 8k), providing an explanation for the broader activation and greater spatial synchrony of auditory neurons responses to acoustic input. Together, these results reveal that although spontaneous activity is preserved in the absence of Cx26, cessation of this burst firing after ear canal opening and the resulting loss of peripheral input results in rapid adaptation of auditory neurons, leading to enhanced gain and broader frequency tuning of neurons within central auditory centers.

## DISCUSSION

Loss of function mutations in *Gjb2* are a leading cause of inherited deafness worldwide, but the mechanisms responsible these hearing deficits are unknown and there are no approved therapies to restore Cx26 function in these patients. Despite evidence of developmental changes in the structure of the organ of Corti and progressive hair cell degeneration^15^, children with *Gjb2* variants respond better to cochlear implants following early implantation relative to other deafness etiologies^16, 58, 59^, suggesting that their central auditory circuits can process acoustic information effectively. Our studies, which involve analysis of mice in which *Gjb2* was selectively inactivated within the cochlear sensory epithelium, reveal that intrinsically generated spontaneous activity that emerges from the developing cochlea prior to hearing onset persists in absence of these gap junctions, providing the means to promote neuronal survival, refine frequency tuning, and establish proper gain of auditory neurons during a critical developmental period^60^. Preservation of this spontaneous activity was unexpected, as cochlear supporting cells form a highly interconnected syncytium using Cx26 gap junctions, which coordinates their maturation and enables ion redistribution following mechanotransduction. Moreover, Cx26 hemichannels can serve as a conduit for the release of ATP^26^, which initiates these periodic bursts of activity in the cochlea prior to hearing onset. Nevertheless, ISCs in the developing cochlea remained highly coupled in the absence of Cx26 and retained the ability to release ATP in a spatially and temporally restricted manner, allowing coordinated excitation of groups of IHCs along the length of the cochlea. These studies highlight the unexpected resilience of both the pan-supporting cell syncytium in the organ of Corti and the mechanisms responsible for initiating spontaneous activity in the developing auditory system. Our studies also reveal that cessation of spontaneous activity and deficiencies in transmission of acoustic information at hearing onset led to a rapid, progressive increase in auditory neuron excitability, insight that may help improve the performance of cochlear implants in patients with mutations in *Gjb2*.

### Intercellular coupling and ATP release in the absence of Cx26

Gap junctions have been implicated in coordinating the development and physiological state of the cochlear sensory epithelium by allowing transfer of microRNAs and second messengers such as IP3 and calcium through the supporting cell syncytium^61–63^. This intercellular signaling is thought to be mediated primarily by Cx26 and Cx30, the most prevalent gap junction proteins expressed by supporting cells in both the developing and mature cochlea^6, 64, 65^. Over 50 distinct mutations in *Gjb2* have been identified and studies in heterologous expression systems indicate that most result in loss of function due to impaired trafficking and assembly^10^. Moreover, genes encoding these proteins, *Gjb2* and *Gjb6*, respectively, are adjacent in the genome and their expression is co-regulated, with mutations in *Gjb2* lowering *Gjb6* transcription and vice-versa^26^, suggesting that disruption of *Gjb2* should substantially impair coupling and coordinated cellular signaling in the organ of Corti. Although tracer spread between ISCs was reduced, and ATP-dependent increases in calcium and resulting osmotic crenations were more spatially restricted in Cx26 cKO mice, supporting cells adjacent to IHCs (including future inner phalangeal cells) in Cx26 cKO mice retained gap junctional coupling sufficient to enable transfer of tracer and high electrical coupling. It is not yet known which gap junctions enable this preservation of coupling or whether it arises through retained expression or compensatory upregulation of latent gap junctions. Recent single-cell sequencing of the developing sensory epithelium suggests that additional gap junctions are expressed within Kölliker’s organ at this age, including *Gja1* (Cx43) and *Gjc1* (Cx45)^66–68^, albeit at substantially lower levels compared to *Gjb2* or *Gjb6*. Although ISCs remained coupled in the absence of Cx26, developmental abnormalities in the structure of the cochlear epithelium remained, including agenesis of the tunnel of Corti and Nuel’s space, demonstrating a distinct requirement for Cx26 beyond ISC intercellular coupling.

Connexin hemichannels (unpaired connexin channels in the plasma membrane) can allow ATP release^26, 69^, raising the possibility that they may enable activation of P2RY1 autoreceptors on ISCs^32, 40^. Despite the high expression of Cx26 in ISCs, spontaneous ATP release events persisted in Cx26 cKO mice, indicating that Cx26 (and Cx30) hemichannels are not essential conduits for ATP release in the developing cochlea. Pannexin channels and purinergic P2rx7 channels can also mediate ATP release and Pannexin 1 is expressed by ISCs^66^; however, spontaneous calcium transients also persist in ISCs of pre-hearing cochlea in the absence of Pannexin 1^70^ or P2rx7^71^channels. It is plausible that remaining gap junction hemichannels or other anion channels enable ATP release. The low frequency of these events suggest that spontaneous activity could be supported by only a few channels per cell if their gating probability and ATP permeability were sufficiently high.

### Early establishment of tonotopic organization and cortical gain in deaf mice

Cx26 cKO mice exhibited profound auditory dysfunction following ear canal opening, with elevated ABR thresholds and reduced activation of neurons in the IC and AC as early as P13. Nevertheless, ABRs and sound-evoked neural calcium increases in the IC and AC could still be elicited with suprathreshold (>80-90 dB SPL) pure tone stimuli at these early ages, indicating that cochlear mechanotransduction and signal propagation through auditory circuits still occurs. Recent studies indicate that pre-hearing spontaneous activity is required to establish proper gain and frequency tuning of central auditory neurons, as well as consolidate sound processing domains within the brain^60^. Although Cx26 cKO mice were less sensitive to airborne sound, evoked calcium responses in IC and AC mirrored the amplitudes, spatial activation, and tonotopic organization of neuronal activity of control mice, suggesting that the persistence of prehearing spontaneous activity promoted restricted IHC activation along the sensory epithelium and refinement of auditory neuron projections during the pre-hearing period. These studies also indicate that alterations in cochlear structure and abnormal physiological maturation of IHCs in Cx26 or Cx30^11, 72^ deficient mice do not prevent IHC burst firing or propagation of tonotopically restricted spontaneous events to the CNS needed to promote some aspects of circuit maturation.

Despite preservation of normative patterns of neural activity prior to ear canal opening, auditory circuits in Cx26 cKO mice rapidly changed after this intrinsically induced activity ceased and IHC activation became dependent on airborne sound. The low efficiency of sensory transduction within the cochlea of Cx26 cKO mice and corresponding decline in activity propagating through central auditory circuits was accompanied by enhanced gain to suprathreshold pure tones. Prior studies have shown that neurons in auditory cortex increase their excitability after hearing loss^73–76^, which is associated with elevated spontaneous cortical activity and enhanced neural synchrony to residual stimulation that often overshoot beyond their baseline levels^77–80^. Our results indicate that these changes in cortical circuits also occur following congenital hearing loss, and that hypersynchrony to residual stimuli emerges primarily as a result of engagement of normally latent cortical neurons, presumably in part due to decreased inhibition^73, 74, 81, 82^. Although attempting to compensate for reduced peripheral input, this increased central gain could prove detrimental and contribute to suprathreshold hyperacusis (disordered loudness perception) and tinnitus (perception of sound in absence of an acoustic stimulus)^83–86^. While the precise mechanisms underlying the emergence of tinnitus remain unclear and are likely multifactorial^85, 87, 88^, cochlear implants have been shown to reduce tinnitus in individuals with severe hearing loss, suggesting that restoration of peripheral input may reverse these homeostatic changes^89, 90^ and that early intervention to restore hearing may improve functional outcomes in forms of congenital deafness arising from mutations in *Gjb2*.

### Mechanisms of peripheral auditory dysfunction due to loss of Cx26

The reduction in acoustic sensitivity observed in Cx26 cKO mice resembles that of cochlear-wide Cx26 deletion^12, 34, 35^, including elevation of ABR thresholds (>50 dB SPL), loss of DPOAEs, and progressive degeneration of hair cells. However, hair cell degeneration was delayed in Cx26 cKO mice relative to other models^12, 34^, allowing assessment of acoustic sensitivity prior to hair cell loss. Remarkably, deficiencies in sound detection manifested in Cx26 cKO mice prior to hair cell degeneration. Several other factors could contribute to this early auditory dysfunction, including structural abnormalities in the organ of Corti and impaired endocochlear potential generation. In particular, Cx26 cKO mice failed to form the tunnel of Corti and Nuel’s space^11, 35^, perhaps due to abnormal maturation of pillar cells and Deiters’ cells. This structural disruption could impair cochlear amplification by preventing efficient coupling of OHC electromotility to the basilar membrane, leading to loss of DPOAEs and elevated ABR thresholds, as observed following OHC degeneration, supporting cell ablation, or compromised electromotility^91, 92^. However, disruption of gap junction coupling specifically within pillar and Deiters’ cells did not affect formation of the tunnel of Corti or Nuel’s space, and reduced DPOAE sensitivity was only observed within basal cochlear regions^93, 94^, although these studies were not designed to assess an early developmental requirement for Cx26 expression and may have been hampered by incomplete *Gjb2* inactivation in *Prox1-CreER* mice^92, 95^.

Endocochlear potential (EP) reductions of 50-70% have been observed in models of both cochlea-wide and organ of Corti restricted deletion of Cx26^11, 34^, which could also impair hair cell mechanotransduction and cochlear amplification. However, mice that express a dominant negative variant of Cx26 (R75W) exhibit auditory dysfunction, hair cell degeneration, and collapse of the tunnel of Corti and perilymphatic spaces, but retain a normal EP^96^, suggesting that structural changes within the sensory epithelium may be sufficient to impair auditory sensitivity. As spontaneous activity in the prehearing cochlea does not require EP generation or OHC function and cochlear amplification, these impairments only manifest when spontaneous bursts of activity in the auditory nerve cease after ear canal opening. This mechanistic independence of intrinsically generated and sound induced activities may enable preservation of early patterned activity in the developing auditory system in individuals that will later present with profound hearing impairment.

## Acknowledgements

We thank members of the Bergles laboratory for discussions and comments on the manuscript. We thank Michele Pucak and Abigail Bush in the Multiphoton Imaging Core and Terry Shelley in the Neuroscience Machine Shop for assistance. Funding was provided by grants from the National Institutes of Health (R01DC008060, P30NS050274) to Dwight E. Bergles and (U19NS107464, R01DC009607) to Patrick O. Kanold. Calvin J. Kersbergen is supported by an individual NRSA fellowship (F30DC018711) and a Medical Scientist Training Program grant (T32GM136577) from the National Institutes of Health. Travis A. Babola is supporting by an individual NRSA fellowship (F32DC019842) from the National Institutes of Health.

## Author Contributions

Calvin J. Kersbergen: Conceptualization, Methodology, Investigation, Formal analysis, Funding acquisition, Writing--original draft; Travis A. Babola: Methodology, Investigation, Formal analysis, Funding acquisition, Writing--review and editing; Patrick Kanold: Tools and reagents, Funding acquisition, Writing--review and editing; Dwight E. Bergles: Conceptualization, Methodology, Supervision, Funding acquisition, Writing--original draft, Writing--review and editing.

## METHODS

Requests for sharing resources, tools, code, and reagents should be directed to the corresponding author, Dwight E. Bergles.

### Animals

This study was performed in strict accordance with the recommendations provided in the Guide for the Care and Use of Laboratory Animals of the National Institutes of Health. All experiments and procedures were approved by the Johns Hopkins Institutional Care and Use Committee (Protocol #M018M330, #M021M290). Generation and genotyping of transgenic mice, including *Cx26^fl/fl^* (European Mouse Mutant Archive EM:00245)^11^, *Tecta-Cre* (JAX Stock No. 035552)^32^, *R26-lsl-GCaMP3*^41^*, R26-lsl-eGFP* (MMRRC Stock No. 32037-JAX)^97^ and *Snap25-T2A-GCaMP6s* (JAX Stock No. 025111)^43^ have been previously described. All animals were backcrossed and maintained on a C57Bl/6N background. Both male and female mice were used for all experiments.

### Cochlea Histology and Immunohistochemistry

Mice were deeply anesthetized with inhaled isoflurane (< P14) or intraperitoneal injection of 10 mg/ml sodium pentobarbital (> P14) and cardiac perfused with ice cold 1X PBS followed by 4% paraformaldehyde (PFA) in 0.1 M phosphate buffer, pH 7.4. The inner ear was isolated from the temporal bone, post-fixed in 4% PFA overnight at 4 °C and stored in 1X PBS with 0.1% sodium azide until processing. For cross sections, cochleae were decalcified in 10% EDTA in 0.1 M phosphate buffer (pH 7.4) shaking at 4 °C (P7: 2-4 hours, >P14: 48 hours), cryopreserved in 30% sucrose, embedded in O.C.T. compound (Tissue Tek), cut in 10 μm sections using a cryostat, and placed on slides (SuperFrost Plus, Fisher). Cochlea sections and free-floating whole cochlea were preincubated in blocking solution (0.3-0.5% Triton X-100, 5% Normal Donkey Serum in PBS, pH 7.4) and incubated at 4 °C with primary antibodies (Mouse anti-Cx26, 1:100, Invitrogen; Rabbit anti-Cx26; 1:250, Invitrogen; Rabbit anti-Cx30, 1:250, Invitrogen; Rabbit anti-MyosinVIIa, 1:300, Proteus Biosciences; Goat anti-Calbindin, 1:200, Santa Cruz, Rabbit anti-β3-tubulin, 1:500, Cell Signaling Technologies). Following overnight incubation, cochleae were washed 3 x 10 minutes in PBS and incubated with corresponding donkey secondary antibodies (Alexa Fluor 488, Alexa Fluor 546 and Alexa Fluor 647; 1:2000, Invitrogen) for 2 hours at room temperature. For assessment of Neurobiotin spread, cochleae were incubated with conjugated Alexa Fluor 555-Streptavidin (1:1000) for 2 hours during secondary antibody incubation. Finally, slides were washed, incubated with 1:10000 DAPI, and sealed using Aqua Polymount (Polysciences, Inc). For hematoxylin and eosin staining, cochleae were decalcified, dehydrated in 70% ethanol, embedded in paraffin, cut in 5 μm sections, and stained by the Reference Histology core in the Department of Pathology at Johns Hopkins Hospital. Images were captured using an epifluorescence microscope (Keyence BZ-X) or a laser scanning confocal microscope (LSM 880, Zeiss).

### Quantification of Hair Cells and Spiral Ganglion Neurons

For analysis of hair cell density, decalcified cochleae were dissected and cut into 3 segments of equal length (apex, middle, base), and stained for hair cell markers as described above. Images of MyoVIIa immunoreactivity were collected at 25x magnification at the mid-portions of each segment corresponding roughly to 8 kHz, 24 kHz, and 50 kHz, respectively. Quantification of cell density was performed in a blinded fashion using a custom cell counting algorithm (MATLAB), described below, or manually, when restricted by low signal-to-noise. For automated counts and analysis of degenerating OHCs, maximum intensity projections of hair cells were thresholded and binarized, a mask placed over the OHC region, and counts of “preserved” OHC centroids were identified using the MATLAB function ‘bwboundaries’.

For analysis of spiral ganglion neurons, 10 µm-thick mid-modiolar cross-sections were identified, and images collected from the apical, middle, and basal spiral ganglia visible within that section. SGN density was determined in a blinded manner by first measuring the spiral ganglion area in ImageJ and subsequent manual counts of all visible Tuj1-labeled neuronal soma. If multiple mid-modiolar sections were visualized on the same slide, images were collected from all regions and averaged.

### Auditory Brainstem Response (ABR) and Distortion Product Otoacoustic Emission (DPOAE) Measurements

For both ABR and DPOAE measurements, mice were anesthetized by intraperitoneal injection of Ketamine (100 mg/kg) and Xylazine (20 mg/kg) and placed in a sound attenuation chamber. Body temperature was maintained at 37 °C using a rectal temperature probe and feedback heating pad (ABRs) or an isothermal heating pad (DPOAEs).

For ABR assessment, subdermal platinum needle electrodes (E2, Grass Technologies) were placed posterior to the pinna, at the vertex, and leg (ground). Stimuli and signal acquisition were controlled using a custom MATLAB program^98^. Acoustic stimuli consisted of 1 ms clicks and 5 ms tone pips of varying frequency (8, 16, 24, and 32 kHz) at a rate of 20 Hz. Signals were generated by a TDT System 3 (Tucker Davis Technologies), amplified (Crown CH1, Crown Audio Inc.), and delivered through a free-field speaker (FD28D, Fostex) placed 30 cm away from the pinna with 5-10 dB attenuation steps. The speaker was calibrated with a free-field microphone (type 4939, Brüel & Kjær) at 30 cm. ABR signals were amplified (Iso-80, World Precision Instruments) band-pass filtered (Krohn-Hite Model 3550, Krohn-Hite Corporation), digitized (RX6 Multifunction Processor, Tucker Davis Technologies), and averaged across 300 stimuli. ABR thresholds were calculated in an automated manner as the lowest stimulus intensity determined by linear interpolation that produced peak-to-peak ABR signals that were greater than 2 standard deviations above the peak-to-peak background signal. Average ABR traces represent mean +/- standard deviation (shaded region) across all animals.

For DPOAE assessment, an ER10B+ microphone probe (Etymotic Research) with a 3 mm eartip (Etymotic Research 10D-T03) was securely placed in the external ear canal. Continuous pure tones (F1 and F2) were generated by an RZ6 Processor (Tucker Davis Technologies) and presented using 2 closed-field MF1 speakers (Tucker Davis Technologies) connected to the microphone probe by 3 cm of 1/8” PVC tubing. Tones were based at 5 center frequencies (Fc: 8, 12, 16, 20, and 24 kHz) with F1 = 0.909*Fc and F2 = 1.09*Fc. Primary tones were presented in 5 dB attenuation steps from 90 dB SPL to 20 dB SPL. Recording was performed at 20 ms intervals, and responses were averaged 300 times per attenuation level. Tone presentation and recording was controlled by BioSigRZ software (Tucker Davis Technologies). All DPOAEs were recorded blinded to animal genotype. DPOAE thresholds were calculated in an automated manner in MATLAB as the lowest stimulus intensity determined by linear interpolation that produced a peak at the expected DPOAE location (2F1-F2) that were greater than 2 standard deviations above the mean background signal adjacent to that frequency location. Average DPOAE traces represent mean +/- standard deviation (shaded region) across all animals of a given genotype.

### Electrophysiology

Apical segments of the cochlea were acutely isolated from P6-P8 mouse pups and used within 2.5 hours. Cochlea were dissected and subsequently superfused by gravity at 2 mL/minute with bicarbonate-buffered aCSF at room temperature (22-24 °C) containing (in mM): 115 NaCl, 6 KCl, 1.3 MgCl_2_, 1.3 CaCl_2_, 1 NaH_2_PO_4_, 26.2 NaHCO_3_, 11 D-glucose saturated with 95% O_2_ / 5% CO_2_ at a pH of 7.4. Whole cell recordings from ISCs were made under visual guidance using differential interference contrast transmitted light. Electrodes had tip resistances between 2.5 and 4.0 MΩ with internal solution of (in mM): 134 KCh_3_SO_3_, 20 HEPES, 10 EGTA, 1 MgCl_2_, 0.2 Na-GTP, pH 7.4. Measurements of membrane resistance in ISCs were obtained following break in through 10 mV voltage steps from −20 to +20 mV relative to the holding potential. Spontaneous currents were recorded with ISCs held at −80 mV for a minimum of 10 minutes. For cell filling experiments, Neurobiotin (Vector Laboratories) was included in the internal solution at a final concentration of 0.2%. Recordings with Neurobiotin in the patch pipette were not included in analysis of spontaneous activity or input resistance, but did not exhibit any significant differences from recordings without Neurobiotin. For recordings from IHCs, electrodes had tip resistances between 4.0 and 6.0 MΩ with internal solution of (in mM) 134 KCh_3_SO_3_, 20 HEPES, 10 EGTA, 1 MgCl_2_, 0.2 NaCl, pH 7.4. Resting potential measurements were made immediately following cell membrane rupture in current clamp configuration with no current injection.

Measurements of membrane resistance in IHCs were obtained through repeated 10 mV voltage steps from −10 to +10 mV relative to the holding potential, and I-V curves were generated through 10 mV voltage steps from −150 to +20 mV. Spontaneous currents were recorded with IHCs held at −70 mV for a minimum of 15 minutes. Recordings were performed using pClamp 9 software with a Multiclamp 700A amplifier (Axon Instruments), low pass Bessel filtered at 1 kHz, and digitized at 5 kHz (Digidata 1322a, Axon Instruments). Recordings exhibiting > 20% change in access resistance or with access resistance > 30 MΩ at the start of recording were discarded. Errors due to voltage drop across the series resistance and liquid junctional potential were left uncompensated. Application of MRS2500 (Tocris, 1 μM) was performed by superfusion by gravity of MRS2500-containing aCSF without interruption of continuous aCSF flow. Analysis of input resistance, resting membrane potential, and spontaneous activity was performed offline in MATLAB. Spontaneous currents were detected using the ‘peakfinder’ function, with a fixed peak threshold (baseline + 3 standard deviations) and minimum peak amplitude (10 pA for ISCs, 5 pA for IHCs). Analysis of changes in spontaneous activity with MRS2500 was performed 1.5 minutes after change in perfusion, to allow time for inflow of drug into the bath.

### Transmitted Light Imaging

For time-lapse imaging of spontaneous osmotic crenations, acutely excised cochleae were visualized using DIC optics through a 40x water-immersion objective coupled to a 1.8x adjustable zoom lens (Zeiss) on a Zeiss Axioskop 2 microscope. Images were acquired at one frame per second using a frame grabber (LG-3; Scion) and Scion Image software. Crenations were detected by generation of difference movies in Matlab through subtraction of frames at time t_n_ and t_n+5_ seconds. To detect transmittance change events, a threshold of the mean + three standard deviation was applied to the difference signal. To calculate area of crenation events, a Gaussian filter (sigma = 12) was applied to the image after thresholding and the borders detected using MATLAB (‘bwlabel’ function). The crenation area was calculated as the number of pixels within the border multiplied by a scaling factor (μm/pixel)^2^.

### Cochlea Explant Culture and Calcium Imaging

Cochlea segments were acutely isolated from P5-P6 mice in ice-cold, sterile filtered, HEPES buffered artificial cerebrospinal fluid (aCSF) containing (in mM): 130 NaCl, 2.5 KCl, 10 HEPES, 1 NaH_2_PO_4_, 1.3 MgCl_2_, 2.5 CaCl_2_, 11 D-glucose, as previously described^22, 42^. Explants were mounted onto Cell-Tak (Corning) treated coverslips and incubated at 37 °C for 12 hours in Dulbecco’s modified Eagle’s medium (F-12/DMEM; Invitrogen) supplemented with 1% fetal bovine serum (FBS) and 10 U/mL penicillin (Sigma) prior to imaging. After overnight culture, cochleae were transferred to recording chamber and superfused with bicarbonate-buffered aCSF containing (in mM): 115 NaCl, 6 KCl, 1.3 MgCl_2_, 1.3 CaCl_2_, 1 NaH_2_PO_4_, 26.2 NaHCO_3_, 11 D-glucose saturated with 95% O_2_ / 5% CO_2_ at a pH of 7.4 at room temperature (22-24 °C). For IHC and ISC imaging, cochleae were illuminated with a 488 nm laser (maximum 25 mW power), and optical sections containing both IHCs and ISCs were obtained with a pinhole set to 3.67 Airy units, corresponding to 5.4 μm of z-depth. For SGN imaging, the pinhole was set to 2.42 Airy units, corresponding to 3.6 μm of z-depth. Addition of NBQX disodium salt (Tocris, 50 μM) in aCSF and high-potassium aCSF (containing (in mM): 80 NaCl, 40 KCl, 1.3 MgCl_2_, 1.3 CaCl_2_, 1 NaH_2_PO_4_, 26.2 NaHCO_3_, 11 D-glucose saturated with 95% O_2_ / 5% CO_2_ at a pH of 7.4) was performed through superfusion by gravity without interruption of continuous aCSF flow. Images were captured at 2 Hz using a using a Zeiss laser scanning confocal microscope (LSM 710, Zeiss) through a 20X objective (Plan APOCHROMAT 20x/1.0 NA) at 512 x 512 pixels (425.1 by 425.1 microns).

### Analysis of *In Vitro* Calcium Imaging

All image analysis was performed in MATLAB. To assess spontaneous calcium activity within ISCs, a grid of 10 x 10 pixel squares were overlaid on the ISC region, as previously described^40^. Images were normalized to the 10^th^ percentile for each pixel (F_o_), and fluorescence changes (ΔF/F_o_, where ΔF = F – F_o_) were calculated for each ROI over the course of the 10 minute movie. Each grid was binarized as active when ΔF/F_o_ was greater than median + 3 standard deviations of the ΔF/F_o_ signal. ISC events were defined by contiguous activation of connected ISC ROIs in 3 dimensions, with removal of events lasting fewer than 3 frames or less than 5 connected ROIs. Temporally and spatially overlapping events were separated by k-means clustering and visually confirmed as distinct calcium transients.

To assess spontaneous calcium activity within IHCs, oval ROIs aligned with the axis of the cell body were placed over each IHC soma within the field of view, with careful exclusion of phalangeal cell processes, and pixels within each ROI were averaged. Fluorescence changes with time were normalized as ΔF/F_o_ values, where F_o_ was the median fluorescence intensity during the 10-minute recording period for each ROI. Signal peaks were detected (‘findpeaks’ function) with fixed peak threshold criteria (median + 3 standard deviations of ΔF/F_o_ signal) and minimum peak amplitude (10% ΔF/F_o_). To determine the number of IHCs activated during a given ISC calcium transient, the number of IHCs with event peaks between the start and end time of each ISC calcium event was counted. If temporal windows for ISC events overlapped, IHCs were assigned to the first ISC event in which they are activated, and not counted in subsequent overlapping events. To generate correlation matrices, pairwise Pearson’s correlation coefficients were determined between every possible IHC pair. Mean correlation coefficients were defined as the average of the 80^th^ percentile correlation coefficient between IHCs in the cochlea prep.

To assess spontaneous activity within SGNs, circular ROIs were drawn around each SGN soma on a maximum projection image. Fluorescence changes with time were normalized as ΔF/F_o_ values, where F_o_ was the median fluorescence intensity during the 10-minute recording period for each ROI. Signal peaks were detected (‘findpeaks’ function) with fixed peak threshold criteria (median + 3 standard deviations of ΔF/F_o_ signal) and minimum peak amplitude (10% ΔF/F_o_). Only neurons that exhibited a ΔF/F_o_ response greater than 10% to 40 mM K^+^ wash were included for analysis. Analysis of changes in spontaneous activity with NBQX was performed 1.5 minutes after change in perfusion, to allow time for inflow of drug into the bath.

### *In Vivo* Imaging of Spontaneous Activity

Installation of early neonatal cranial windows has been previously described^22^. Briefly, mice were anesthetized in inhaled isoflurane (4% induction, 1.5% maintenance), the dorsal skull exposed to allow for headbar implantation, and a cranial window placed over the resected intraparietal bone overlying the midbrain or over auditory cortex (3.5 mm lateral to lambda), as previously described. In a subset of experiments, aCSF containing Sulfarhodamine 101 (SR101, 10 μM) was washed over the brain surface before sealing of the cranial window to label astrocytes to enable image registration and motion correction. After 1 hour of post-surgical recovery from anesthesia, neonatal mice were moved into a 15 mL conical tube (to swaddle and limit movement) and head-fixed under the imaging microscope. During imaging, pups were maintained at 37 °C using a heating pad and temperature controller (TC-1000; CWE). Wide field epifluorescence images were captured at 10 Hz using a Hamamatsu ORCA-Flash4.0 LT digital CMOS camera attached to a Zeiss Axio Zoom.V16 stereo zoom microscope illuminated continuously with a metal halide lamp (Zeiss Illuminator HXP 200C). Each recording of spontaneous activity consisted of uninterrupted acquisition over 10 minutes. Two-photon imaging was performed using a Zeiss 710 LSM microscope with two-photon excitation achieved by a Ti:sapphire laser (Chameleon Ultra II; Coherent) tuned to 920 nm. Images were collected at 4 Hz (256 x 256 pixels, 425 x 425 μm) from 150 μm Z-depth in the central IC for a minimum of 10 minutes.

For widefield imaging analysis, images were imported into MATLAB, underwent photobleaching correction by fitting a single exponential to the fluorescence decay and subtracting this component from the signal, and intensities were normalized as ΔF/F_o_ values, where ΔF = F – F_o_ and F_o_ was defined as the tenth percentile value for each pixel. Oval regions of interest were placed over the right and left IC, and signal peaks were identified using built-in peak detection (‘findpeaks’) with a fixed threshold (1% ΔF/F_o_) and minimum peak amplitude (1% ΔF/F_o_). Occasional spontaneous broad activation of the cortex and SC would elicit fluorescence increases in the IC, and these events were not included in the analysis, as they do not appear to be derived from the auditory periphery^22^. For analysis of spatial band width, a 25 x 100 rectangular ROI rotated 45-55 degrees was placed over each inferior colliculus aligned with the future tonotopic axis of the IC. The rectangle was averaged along the short axis, creating a 100 x 1 pixel line scan of the tonotopic axis of the IC for the duration of the time series.

Events were detected using the function ‘imregionmax’ with a fixed threshold (2% ΔF/F_o_). The spatial integral of spontaneous events along the tonotopic axis was estimated using the function ‘trapz’. Line scans of individual events were normalized to the maximum ΔF/F_o_ at that time point, and the band width calculated as the length along the tonotopic axis above the 75^th^ percentile of the peak ΔF/F_o_. Two-photon imaging analysis was performed similarly, with a 50 x 300 pixel rectangular ROI rotated 45-55 degrees was placed over the imaging plane to be perpendicular to spontaneous bands. The rectangle was averaged along the short axis, creating a 300 x 1 pixel line scan of the tonotopic axis of the IC for the duration of the time series. Events were detected using the function ‘imregionmax’ with a fixed threshold (2% ΔF/F_o_). Line scans of individual detected events were normalized to the maximum ΔF/F_o_ at that time point, and the band width calculated as the length along the tonotopic axis above the 75^th^ percentile of the peak ΔF/F_o_.

### Sound-Evoked Calcium Imaging

Cranial windows over inferior colliculus in P13-P15 mice were performed as previously described for P7 animals, with animals allowed to recover a minimum of 2 hours prior to imaging. Auditory cortex windows in P13-P21 mice were installed using a microblade and microscissors to remove a ∼4 mm circular region of skull overlying the right auditory cortex. Following skull removal, the dura was carefully removed using microscissors and exposed brain was continuously immersed in aCSF. A 5 mm coverslip was attached using superglue, and animals were allowed to recover a minimum of 2 hours prior to imaging. During imaging, awake animals were head-fixed but on a freely-rotating tennis ball to simulate natural movement.

Acoustic stimuli for widefield imaging were presented using a free-field speaker (MF1; Tucker Davis Technologies) placed 10 cm from the left ear within a custom sound attenuation chamber with external noise attenuation of 40 dB^22^. Stimuli consisted of 4-6 repetitions of sinusoidal amplitude modulated pure tones (1 s, 10 Hz modulation) from 3 to 48 kHz in ¼ octave intervals. All stimuli were cosine-squared gated (5 ms) and played in a random order at 5 s intervals. Tones were generated within the RPvdsEx software (Tucker Davis Technologies), triggered using the microscope’s frame out signal, and delivered through the RZ6 Audio Processor (Tucker-Davis Technologies). Calibration of the free-field speaker was performed using an ACO Pacific microphone (7017) and preamplifier (4016). Given the flat intensity profile (peak 100 dB SPL +/- 5 dB for a 2.0 V stimulus across all tested frequencies), levels were not corrected across presented frequencies. Stimuli were presented from 100 dB to 20 dB SPL in 10-20 dB attenuation steps.

For sound-evoked two photon imaging, auditory cortex was first identified using widefield imaging, as described above, using a 3-48 kHz tone series at 80-90 dB SPL. The resulting widefield map was used for localization of low and high frequency borders of A1. For two photon imaging, mice were head fixed using a custom 3D-printed headbar and placed on a rotating tennis ball under a Bruker Ultima 2P+ two photon microscope. A 16X Nikon objective (0.8 NA) oriented at 55-60 degrees was immersed in ultrasound gel (ParkerLabs Aquasonic CLEAR) and focused to capture low and high frequency regions of layer 2/3 of primary auditory cortex (250 μm deep from cortical surface) using the widefield mapping and local vasculature as a guide. Two-photon excitation was generated by a Chameleon Discovery (Coherent) laser at 920 nm. Precise imaging location was verified by generating a live “Quick Map” of averaged tone evoked responses using 3 frequency stimuli (4, 16, 64 kHz) presented with 10 repeats at 80 dB SPL, and any adjustments of the imaging field were made to capture both low and high frequency regions of A1 before full dataset acquisition. Acoustic stimulation was generated by an RX6 processor (Tucker Davis Technologies), with 300 ms sinusoidally amplitude-modulated pure tones (4-64 kHz, 9 frequencies in ½ octave increments) triggered every 3 seconds by pre-specified frame out signals. Tones were presented in a random order at 30-90 dB SPL in 20 dB increments through an electrostatic speaker (ES-1, Tucker Davis Technologies). Images were collected using Prairie View software at 1024×1024 pixels (1104 by 1104 microns) at 15 Hz using an 8 kHz resonant galvanometer.

For analysis of sound-evoked widefield responses, raw widefield epifluorescence images underwent bleach correction and normalization as described above for widefield imaging of spontaneous activity. Image segments were separated by tone frequency, aligned from 1 second prior to and 3 seconds following tone presentation, and averaged across the 4 presentations of each tone. Normalized and averaged images were used for display purposes. For analysis of evoked amplitude and auditory thresholds in inferior colliculus, an oval ROI was placed over the contralateral IC. For assessment of spatial band width and frequency mapping, a maximum-intensity projection of the mean sound-evoked response for each presented frequency was rotated by 45-55 degrees and a 25×250 pixel rectangle was placed along the tonotopic axis, centered at the 3 kHz response location or medial IC location for each animal. To generate the sound-evoked spatial profile, ΔF/F_o_ signal intensity was averaged along the short rectangle axis. Profiles were normalized to the maximum ΔF/F_o_ intensity (peak response location) along the tonotopic axis. The normalized spatial band width of the evoked response was defined as the width of the 75^th^ percentile of the normalized response for each mouse. For analysis of auditory cortex responses, a circular ROI was placed over A1 after visualization of averaged pure tone responses. The low and high frequency borders of A1 were defined by their caudal and dorsal locations in auditory cortex^48, 50^. For calculation of pure tone activation area, normalized images were binarized at a fixed threshold (15% ΔF/F_o_) and the total activated area was calculated. Only mice with windows in which the entire auditory cortex was visible were used for analysis. If motion correction was required of widefield movies, it was performed using the MOCO^99^ fast motion correction plugin (ImageJ).

For analysis of two photon sound-evoked calcium imaging, registration, motion correction and soma ROI identification was performed using Suite2P software^100^ and soma and neuropil fluorescence were extracted and imported into MATLAB. Neuropil fluorescence was subtracted from the soma fluorescence (soma – 0.7 x neuropil) and signals were normalized to their mean baseline values (15 frames (1 s) prior to tone stimulus). Data was unmixed relative to the stimulus presented and averaged to generate mean response traces for all frequency and sound intensity stimuli. Principal component analysis was performed on the peak responses following tone stimuli for all frequency and sound intensity combinations, with the three major principal components plotted to demonstrate the distinct response patterns of control and Cx26 cKO cells. Best frequencies were identified as the frequency that elicited the largest calcium response at any sound level. Cells were sorted by their best frequency, and mean evoked responses for each best frequency were plotted at 90 dB SPL. To calculate bandwidth, responsive cells were identified and a single term Gaussian fit (‘fit’, ‘Gauss1’) applied to the maximum evoked amplitudes at each frequency at 90 dB SPL. Cells were excluded if a Gaussian fit could not be made from the maximum amplitudes.

## Statistical Analysis

All statistical testing was performed in MATLAB. All data is presented as mean +/- standard deviation, unless otherwise noted. Datasets were tested for normality using the Lilliefors test (‘lillietest’). If unable to reject the null hypothesis that the dataset is normally distributed using the Lilliefors test, a paired (‘ttest’) or unpaired (‘ttest2’) two-tailed t-test was used to compare groups. If the null hypothesis of normality was rejected, a nonparametric Wilcoxon rank sum (‘ranksum’) or Wilcoxon sign rank (‘signrank’) test was used for unpaired or paired samples, respectively. If multiple comparisons on the same datasets were made, a Benjamini-Hochberg correction of the false discovery rate (‘fdr_BH’) was made to adjust p values to lower the probability of type 1 errors. Comparison of cumulative distributions was performed using a two-sample Kolmogorov-Smirnov test (‘kstest2’). Comparison of multiple datasets of independent samples was made using a one way ANOVA (‘anova1’) with Tukey’s post hoc comparisons. For datasets with multiple comparisons of non-independent samples, a repeated measures ANOVA (‘ranova’) or a linear mixed-effects model (‘fitlme’) were used, using the reduced maximum likelihood fit method (‘FitMethod’,’reml’) and the Satterthwaite approximation of degrees of freedom. Use of a linear mixed model enabled accounting for data dependency for repeated measurements (such as pure tone responses or ABR recordings) from the same mouse as a random effect. Sidák post hoc test was used to assess for post hoc comparisons as indicated in figure legends. Adjusted p-values are displayed as follows: * = p < 0.05, ** = p < 0.01, *** = p < 0.001, **** = p < 0.0001, ns = not significant. Details about number of data points, exact p values, and individual statistical tests can be found in the figure legends.

## EXTENDED DATA FIGURE LEGENDS

**Extended data Figure 1. Morphological anomalies in the organ of Corti with loss of supporting cell Cx26** a) (left) Hematoxylin and eosin stain of mid-modiolar section of P21 control (*Gjb2^fl/fl^*) cochlea. Black square indicates site of high magnification. (right) Magnified image of the organ of Corti, with the tunnel of Corti indicated by black line. (bottom) Schematic depicting morphology of hair cells and supporting cells in the control organ of Corti. IPhC: inner phalangeal cell; IPC: inner pillar cell; OPC: outer pillar cell; DCs: Deiters’ cells; OHCs: outer hair cells; IHC: inner hair cell. b) (left) Hematoxylin and eosin stain of mid-modiolar section of P21 Cx26 cKO (*Tecta-Cre;Gjb2^fl/fl^*) cochlea. Black square indicates site of high magnification. (right) Magnified image of the organ of Corti, with site of the tunnel of Corti indicated by black line. (bottom) Schematic depicting morphology of hair cells and supporting cells in the Cx26 cKO organ of Corti.

**Extended data Figure 2. Hair cell degeneration is progressive with increasing age following loss of Cx26** a) Representative images of hair cells labeled by immunoreactivity to Myosin VIIa (magenta) in whole mounts from apex, middle, and basal regions in control (*Gjb2^fl/fl^*) cochleae at P7, P15, P21, P45, and P100. b) Representative images of hair cells labeled by immunoreactivity to Myosin VIIa (magenta) in whole mounts from apex, middle, and basal cochlea regions in Cx26 cKO (*Tecta-Cre;Gjb2^fl/fl^*) cochleae at P7, P15, P21, P45, and P100. c) Quantification of outer hair cells (OHCs, left) and inner hair cells (IHCs, right) with increasing age in apical (squares), middle (diamonds), and basal (circles) regions of control and Cx26 cKO cochlea.

**Extended data Figure 3. Loss of outer hair cells in Cx26 cKO mice is restricted to basal cochlea at P21** a) Representative images of hair cells labeled by immunoreactivity to Myosin VIIa (magenta) in whole mounts from apex, middle, and basal cochlea in control (*Gjb2^fl/fl^*, left) and Cx26 cKO (*Tecta-Cre;Gjb2^fl/fl^*, right) mice at P21. White arrows indicate sites of outer hair cell loss. b) Quantification of inner hair cells (IHCs, left) and outer hair cells (OHCs, right) at P21 in control and Cx26 cKO apical, middle, and basal cochlea segments. n = 8 control cochleae, 8 Cx26 cKO cochleae; p = 0.0974, 0.0021 (IHCs, OHCs), linear mixed model with Sidák post hoc test. c) (left) Low magnification image of spiral ganglion neurons (SGNs) labeled by immunoreactivity to Tuj1 (cyan) in mid-modiolar cross section of P21 cochlea. Labels indicate locations of apical, middle, and basal SGN counts. (right) Representative high-magnification images of SGN soma labeled by immunoreactivity to Tuj1 in apical, middle, and basal cochlea from control and Cx26 cKO mice at P21. Dashed lines indicate SGN compartment used for area measurement. d) Quantification of SGN density in apical, middle, and basal cochlea at P21. n = 3 control cochleae, 3 Cx26 cKO cochleae; p = 0.4385, repeated measures ANOVA with lower bound p value adjustment.

**Extended data Figure 4. Loss of Connexin 26 in pre-hearing supporting cells reduces osmotic crenation size** a) Immunostaining for Connexin 26 (green) in whole mount middle P7 cochlea from control (*Gjb2^fl/fl^*) and Cx26 cKO (*Tecta-Cre;Gjb2^fl/fl^*) mice. Hair cells (magenta) are labeled by immunoreactivity to Myosin VIIA. b) Immunostaining for Connexin 26 (green) in mid-turn cochlea cross section at P7. Loss of Cx26 immunostaining is observed in Kölliker’s organ and supporting cells of the organ of Corti (white arrows) but not within lateral wall or spiral limbus fibrocytes or the stria vascularis (green arrows). Hair cells are labeled by antibodies against Calbindin (magenta). c) Intrinsic optical imaging of osmotic crenations in control and Cx26 cKO cochleae. Detected crenations are indicated with transparent colored areas based on time of occurrence. d) Quantification of spontaneous crenation area. n = 9 control, 9 Cx26 cKO cochleae; p = 4.4669e-4, two-sample t-test with unequal variances.

**Extended data Figure 5. No changes in electrophysiologic properties of immature inner hair cells with loss of Cx26** a) Schematic of whole cell patch clamp recording from inner hair cells (IHCs). b) Voltage protocol (left) and representative current responses (right) from P7 control (*Gjb2^fl/fl^*) and Cx26 cKO (*Tecta-Cre;Gjb2^fl/fl^*) IHCs. c) Quantification of membrane resistance and resting membrane potential in P7 control and Cx26 cKO IHCs. n = 13 control, 19 Cx26 cKO; p = 0.9838, 0.9847 (membrane potential, membrane resistance), two-sample t-test (membrane potential) or Wilcoxon rank sum test (membrane resistance) with Benjamini-Hochberg correction.

**Extended data Figure 6. Coordinated activation of future isofrequency lamina in pre-hearing Cx26 cKO mice** a) Representative spontaneous neural calcium transients (left) and corresponding normalized spatial fluorescence profile along future tonotopic axis (right, indicated by white rectangle) in IC from P7 control (*Gjb2^fl/fl^;Snap25-T2A-GCaMP6s*) mouse. Merged pseudocolored image (bottom) highlights spatial segregation of discrete spontaneous calcium transients. Band width is calculated as the 75^th^ percentile of the normalized spatial profile. b) Same as (a), but from P7 Cx26 cKO (*Tecta-Cre;Gjb2^fl/fl^;Snap25-T2A-GCaMP6s*) mouse. c) Cumulative distribution of spontaneous event peak location along tonotopic axis of the IC. Gray shading represents central tonotopic region (40^th^ - 60^th^ percentile) used for calculation of band width in (e). n = 1290 control events (7 mice), 1385 Cx26 cKO events (8 mice); p = 0.4908, two-sample Kolmogorov-Smirnov test. d) Quantification of spatial integral of all spontaneous events along tonotopic axis. n = 1290 control events (7 mice), 1385 Cx26 cKO events (8 mice); p = 1.0825e-9, two-sample Kolmogorov-Smirnov test. e) (left) Quantification of mean normalized band width (75^th^ percentile) of spontaneous events along tonotopic axis within central IC. n = 7 control mice, 8 Cx26 cKO mice; p = 0.6620, Wilcoxon rank sum test. f) Representative spontaneous calcium transients in neurons and neuropil within the inferior colliculus of P7 control (left) and P7 Cx26 cKO (right) mice using two-photon imaging. SR101 (magenta) labels astrocytes for image registration. Normalized fluorescence profile for each event was calculated g) (left) Cumulative distribution of normalized band width of spontaneous calcium transients measured using two-photon imaging. n = 236 control events (5 colliculi, 3 mice), 253 Cx26 cKO events (5 colliculi, 3 mice); p = 0.0059, two-sample Kolmogorov-Smirnov test. (right) Quantification of mean band width. N = 5 control colliculi (3 mice), 5 Cx26 cKO colliculi (3 mice); p = 0.0958, two-sample t-test with unequal variances.

**Extended data Figure 7. Topographic organization and response amplitudes to tonal stimuli in developing Cx26 cKO mice** a) Tone-evoked neural calcium transients in IC from P14 control (*Gjb2^fl/fl^;Snap25-T2A-GCaMP6s*, left) and P14 Cx26 cKO (*Tecta-Cre;Gjb2^fl/fl^;Snap25-T2A-GCaMP6s*, right) mice at 100 dB sound pressure level (SPL). Rectangular ROIs were placed along the tonotopic axis of the contralateral IC (low (L) to high (H) frequency), perpendicular to pure tone evoked bands, to determine the peak response location for a pure tone along the tonotopic axis. b) Quantification of peak response location of pure tones along the tonotopic axis relative to the medial IC. n = 4 control mice, 5 Cx26 cKO mice; mean ± SEM, p = 0.0205, linear mixed effects model. c) Quantification of peak response location of pure tones along the tonotopic axis relative to 3 kHz (lowest frequency) peak response location. n = 4 control mice, 5 Cx26 cKO mice; mean ± SEM, p = 0.5654, linear mixed effects model. d) Quantification of tone-evoked fluorescence changes in IC within whole IC ROIs across a range of frequency and sound level stimuli in individual P13-P15 control mice. Each black line indicates average responses from the IC contralateral to acoustic stimulation within an individual animal. Vertical gray bar indicates tone presentation. e) Quantification of tone-evoked fluorescence in IC across a range of frequency and sound level stimuli in individual P13-P15 Cx26 cKO mice. Each red line indicates average responses from the IC contralateral to acoustic stimulation within an individual animal. Vertical gray bar indicates tone presentation.

**Extended data Figure 8. Auditory dysfunction precedes substantial hair cell degeneration** a) Average auditory brainstem response (ABR) traces to broadband click (left) and 16 kHz tone pip stimuli at multiple sound pressure levels from control (*Gjb2^fl/fl^*, black, n = 3) and Cx26 cKO (*Tecta-Cre;Gjb2^fl/fl^*, red, n = 4) at P15-P16. Shaded region represents standard deviation of the average signals across animals. b) Quantification of P15-P16 ABR thresholds to click and pure tone stimuli in controls (*Gjb2^fl/fl^*, black, n = 3 and *Tecta-Cre;Gjb2^fl/+^*, blue, n = 5) and Cx26 cKO (*Tecta-Cre;Gjb2^fl/fl^*, red, n = 4). Detection limit dashed line indicates maximum output from speaker. p = 4.4717e-9 (cKO vs. control), 8.6968e-6 (cKO vs. Cre+ control), 0.1454 (control vs. Cre+ control), linear mixed effects model. c) Representative images of hair cells labeled by immunoreactivity to Myosin VIIa (magenta) in whole mounts from apex, middle, and basal cochlea in control (*Gjb2^fl/fl^*, left) and Cx26 cKO (*Tecta-Cre;Gjb2^fl/fl^*, right) at P15. White arrows indicate sites of outer hair cell loss. d) Quantification of inner hair cells (IHCs, left) and outer hair cells (OHCs, right) at P15-P16 in control and Cx26 cKO apical, middle, and basal cochlea segments. n = 9 control, 9 Cx26 cKO cochleae; p = 0.0326, 0.0053 (IHCs, OHCs), linear mixed model with Sidák post hoc test. e) (left) Hematoxylin and eosin stain of mid-modiolar section of P15 control cochlea. Black square indicates site of high magnification. (right) Magnified image of the organ of Corti, with the tunnel of Corti indicated by black line. f) (left) Hematoxylin and eosin stain of mid-modiolar section of P15 Cx26 cKO cochlea. Black square indicates site of high magnification. (right) Magnified image of the organ of Corti, with site of the tunnel of Corti indicated by black line.

## Supplementary Video Legends

**Supplementary Video 1. Inner supporting cells generate spontaneous activity despite absence of Cx26** Confocal imaging of calcium transients in inner hair cells and inner supporting cells (ISCs) in P7 control (*Tecta-Cre;Gjb2^fl/+^;R26-lsl-GCaMP3*, top) and Cx26 cKO (*Tecta-Cre;Gjb2^fl/fl^;R26-lsl-GCaMP3*, bottom) excised cochlea expressing the genetically encoded calcium indicator GCaMP3. Images were collected at 2 Hz, playback is 10 frames per second.

**Supplementary Video 2. Spontaneous activity in spiral ganglion neurons is mediated by synaptic glutamate release** Confocal imaging of spontaneous calcium transients in spiral ganglion neurons (SGNs) from an isolated Cx26 cKO (*Tecta-Cre;Gjb2^fl/fl^;Snap25-T2A-GCaMP6s*) cochlea. NBQX (50 μM) application eliminates all spontaneous calcium transients. Images were collected at 2 Hz, playback is 20 frames per second.

**Supplementary Video 3. Preserved spontaneous neural activity in the auditory midbrain of Cx26 cKO mice** Wide-field imaging of spontaneous neural activity in the inferior colliculus of P7 control (*Gjb2^fl/fl^;Snap25-T2A-GCaMP6s*, top) and Cx26 cKO (*Tecta-Cre;Gjb2^fl/fl^;Snap25-T2A-GCaMP6s*, bottom) mice that expressed GCaMP6s pan-neuronally under the *Snap25* promoter. Images were collected at 10 Hz, playback is 20 frames per second.

**Supplementary Video 4. Tonotopic organization of suprathreshold pure tone responses in Cx26 cKO mice** Wide-field imaging of sound-evoked neural activity in the auditory cortex of P14 control (*Gjb2^fl/fl^;Snap25-T2A-GCaMP6s*, top) and Cx26 cKO (*Tecta-Cre;Gjb2^fl/fl^;Snap25-T2A-GCaMP6s*, bottom) mice. 3, 12, and 48 kHz sinusoidally amplitude-modulated pure tones are presented for 1 s at 100 dB SPL. Images were collected at 10 Hz, playback is 3 frames per second.

**Supplementary Video 5. Increased gain of evoked responses after hearing onset in Cx26 cKO mice** Wide-field imaging of sound-evoked neural activity in the auditory cortex of P21 control (*Gjb2^fl/fl^;Snap25-T2A-GCaMP6s*, left) and Cx26 cKO (*Tecta-Cre;Gjb2^fl/fl^;Snap25-T2A-GCaMP6s*, right) mice. 3, 16, and 48 kHz sinusoidally amplitude-modulated pure tones are presented for 1 s at 100 dB SPL. Images were collected at 10 Hz, playback is 3 frames per second.

**Supplementary Video 6. Increased engagement of cortical neurons to suprathreshold acoustic stimuli in Cx26 cKO mice** Two-photon imaging of sound-evoked neural activity in the auditory cortex of P21 control (*Gjb2^fl/fl^;Snap25-T2A-GCaMP6s*, left) and Cx26 cKO (*Tecta-Cre;Gjb2^fl/fl^;Snap25-T2A-GCaMP6s*, right) mice. 3, 12, and 48 kHz sinusoidally amplitude-modulated pure tones are presented for 0.3 s at 90 dB SPL. Images were collected at 15 Hz, playback is 4 frames per second.

