## Supplementary Figures for "Preservation of prehearing spontaneous activity enables early auditory system development in deaf mice"

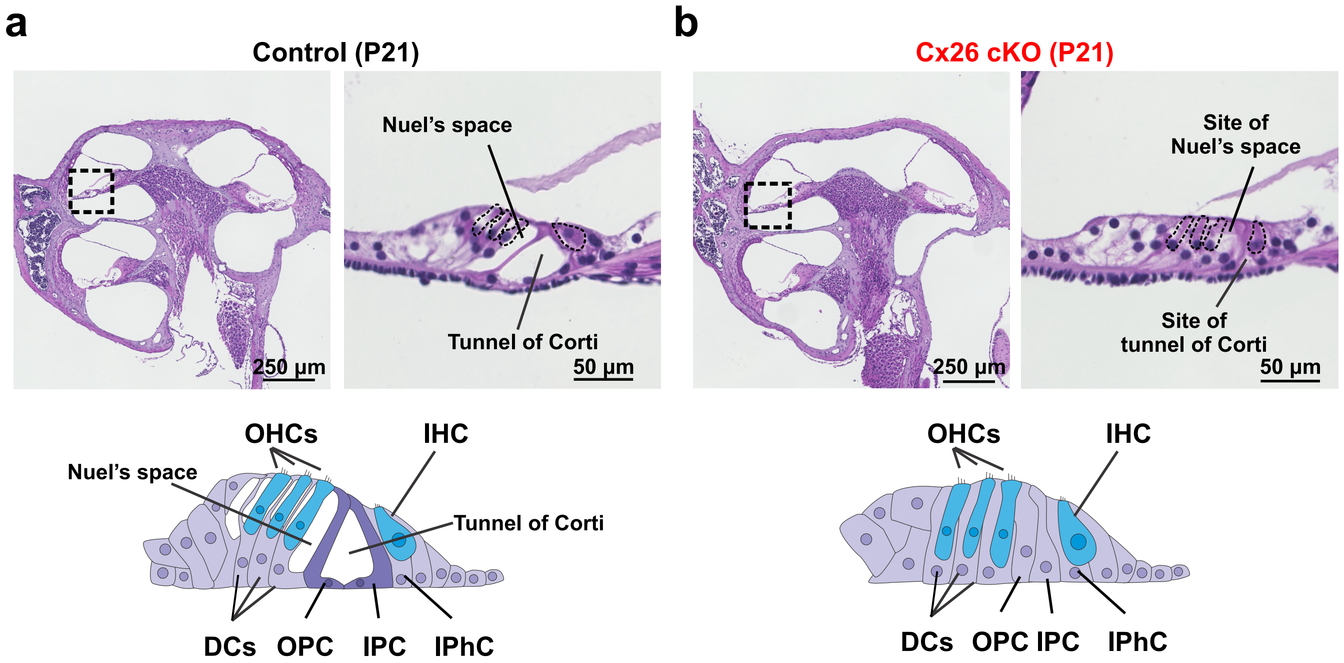


### Extended data Figure 1

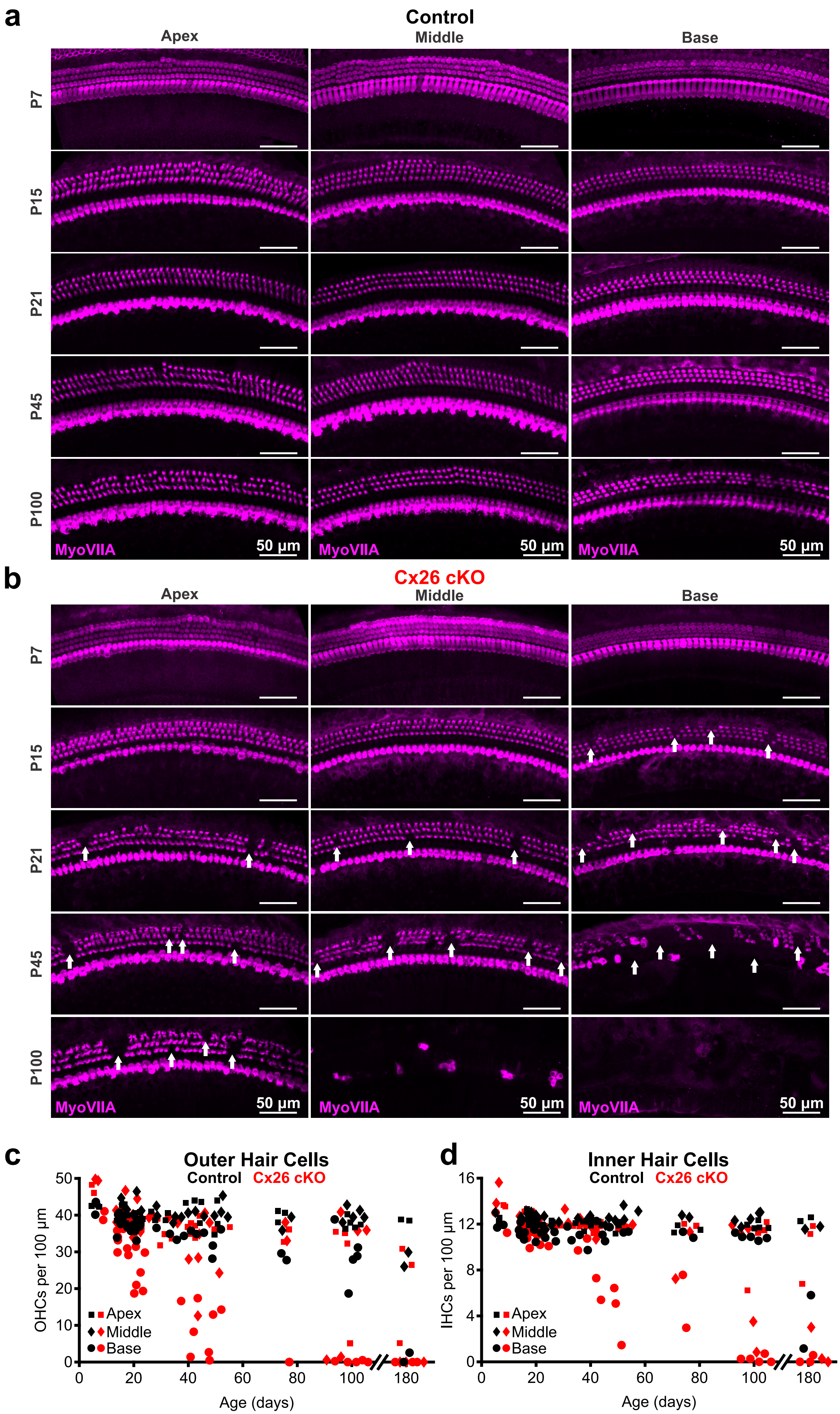


### Extended data Figure 2

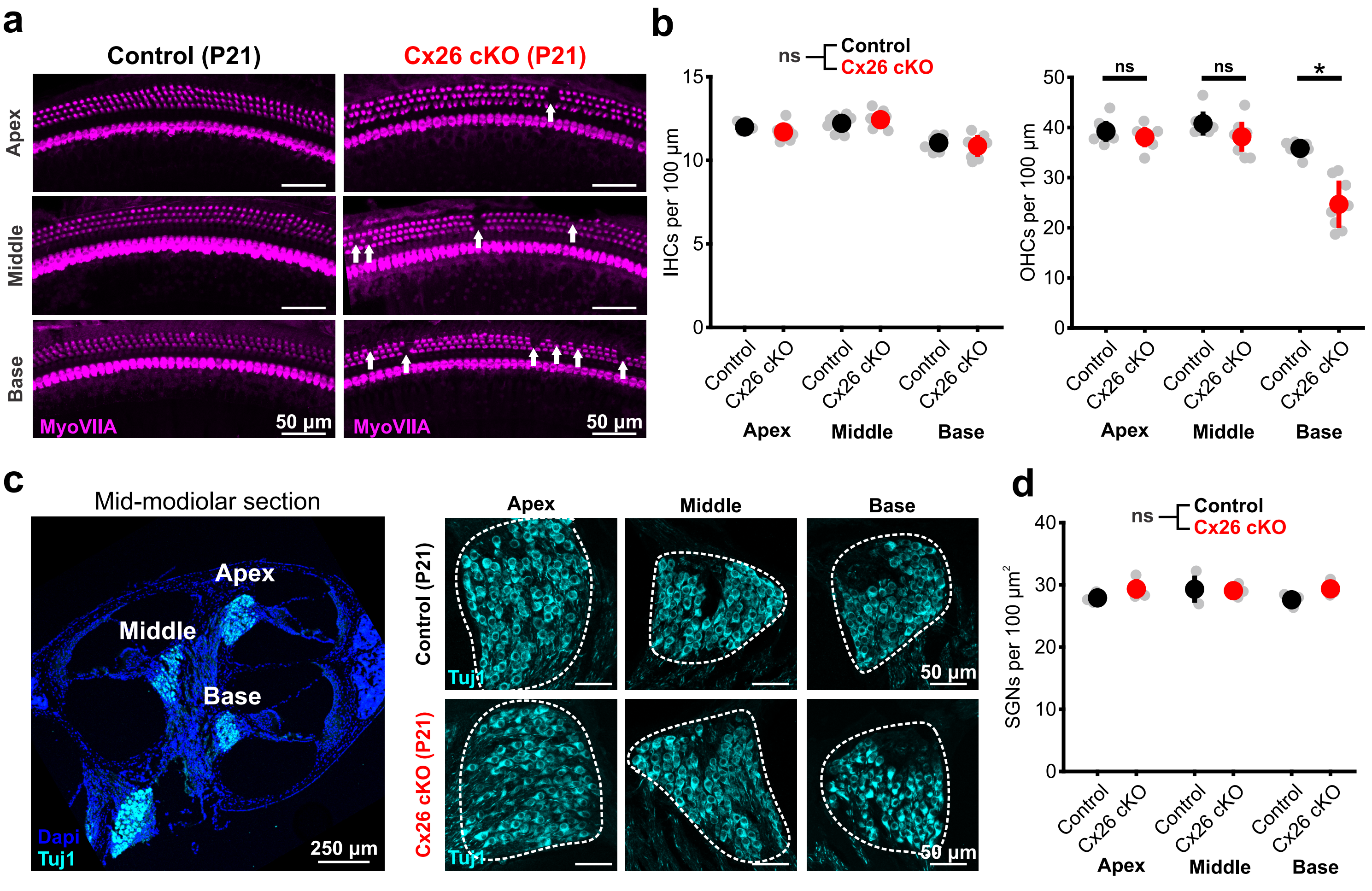


### Extended data Figure 3

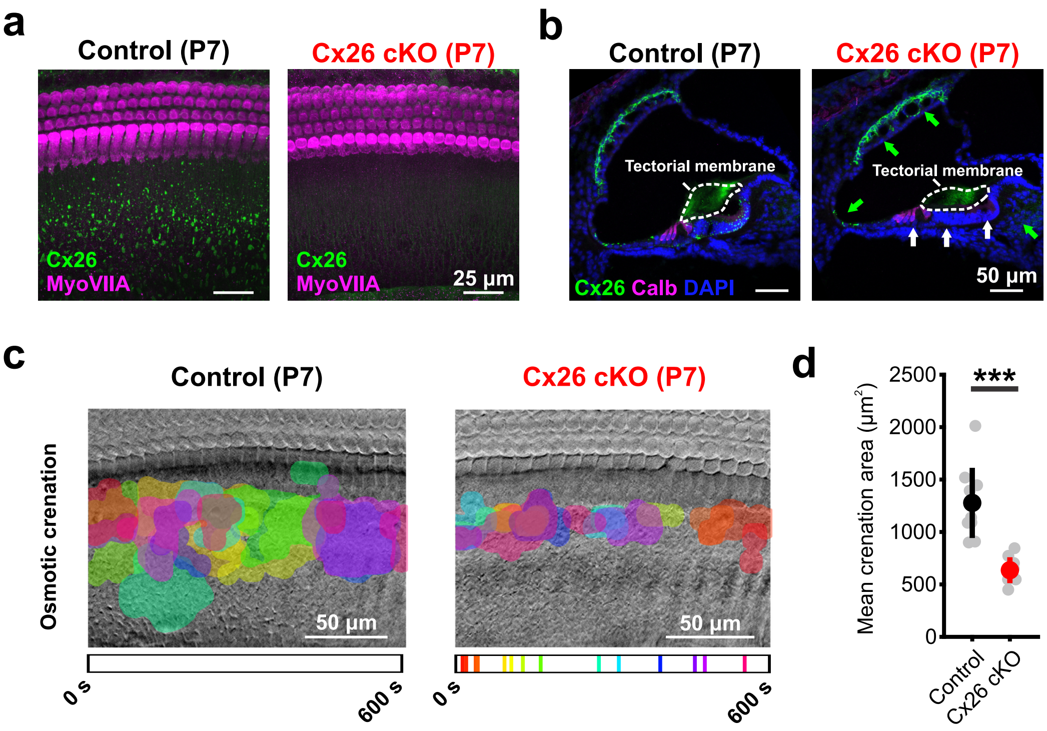


### Extended data Figure 4

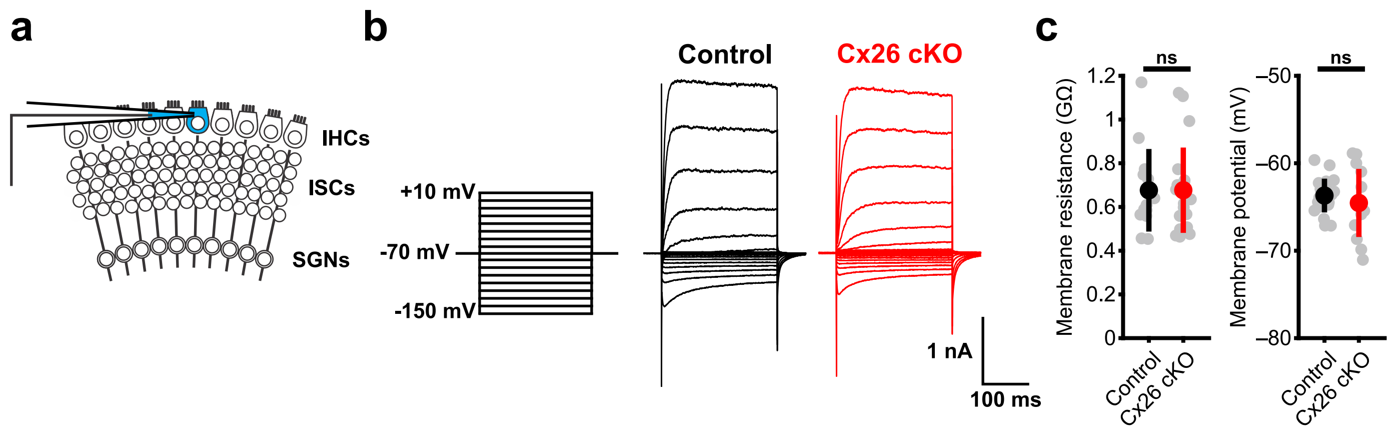


### Extended data Figure 5

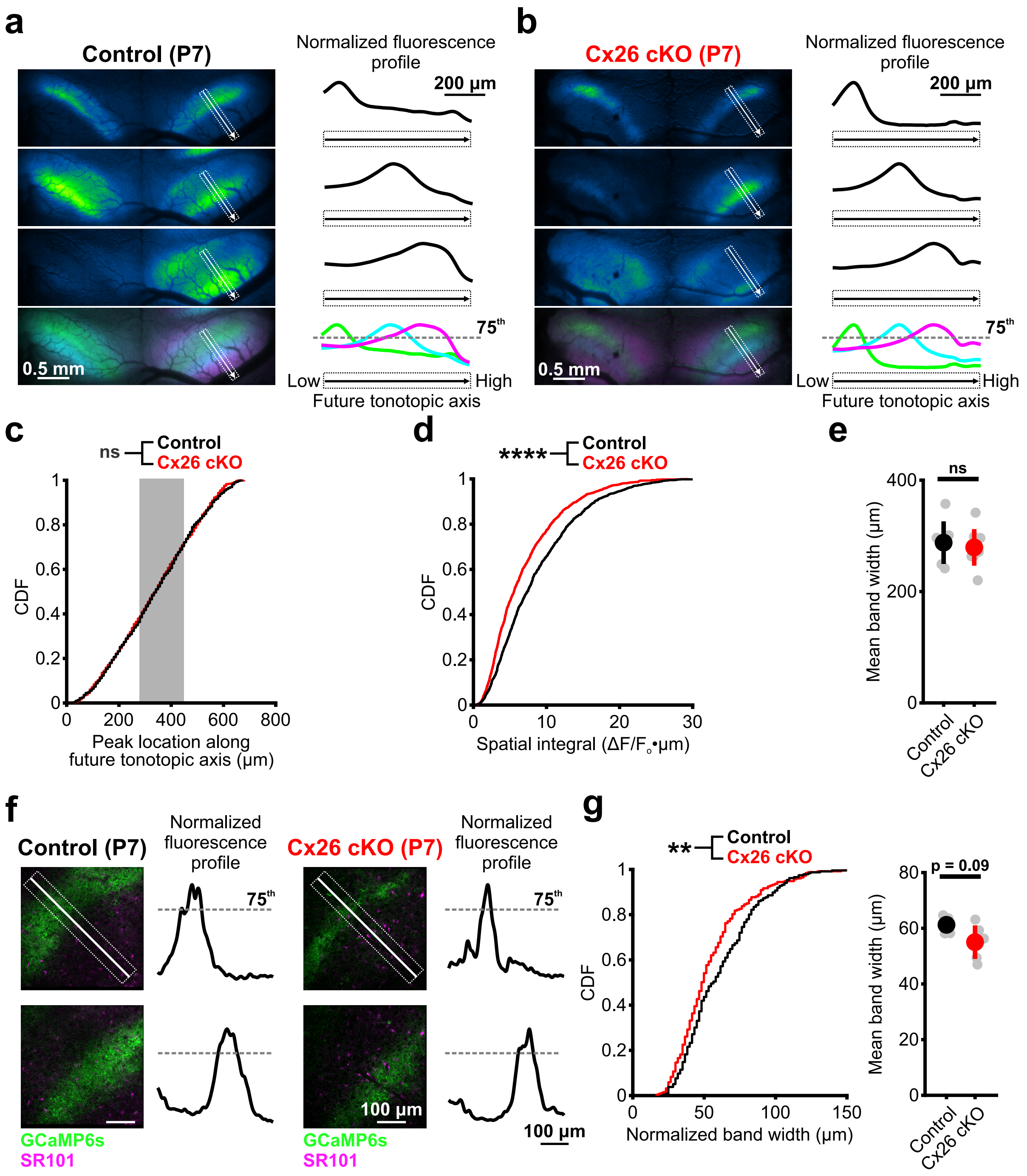


### Extended data Figure 6

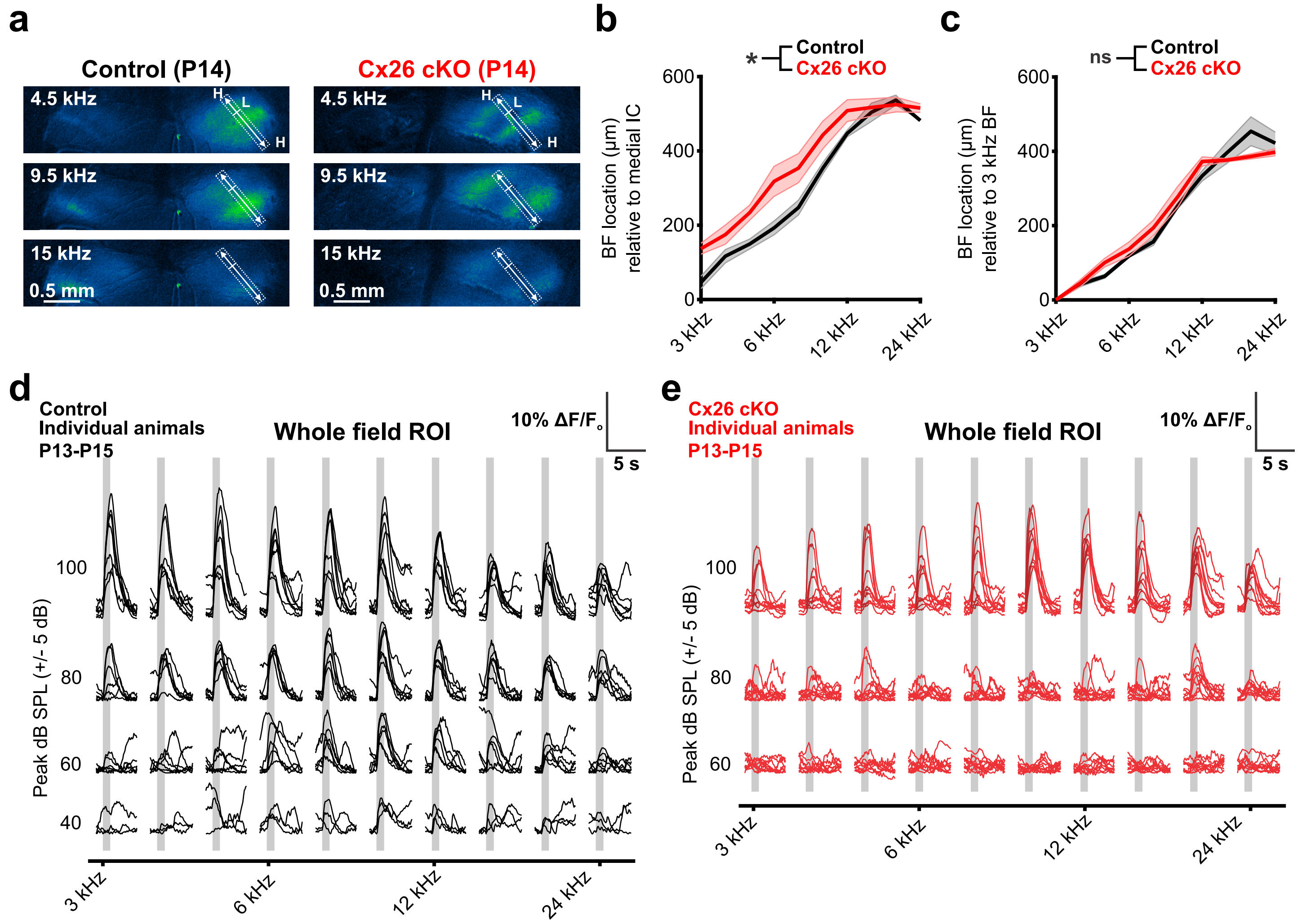


### Extended data Figure 7

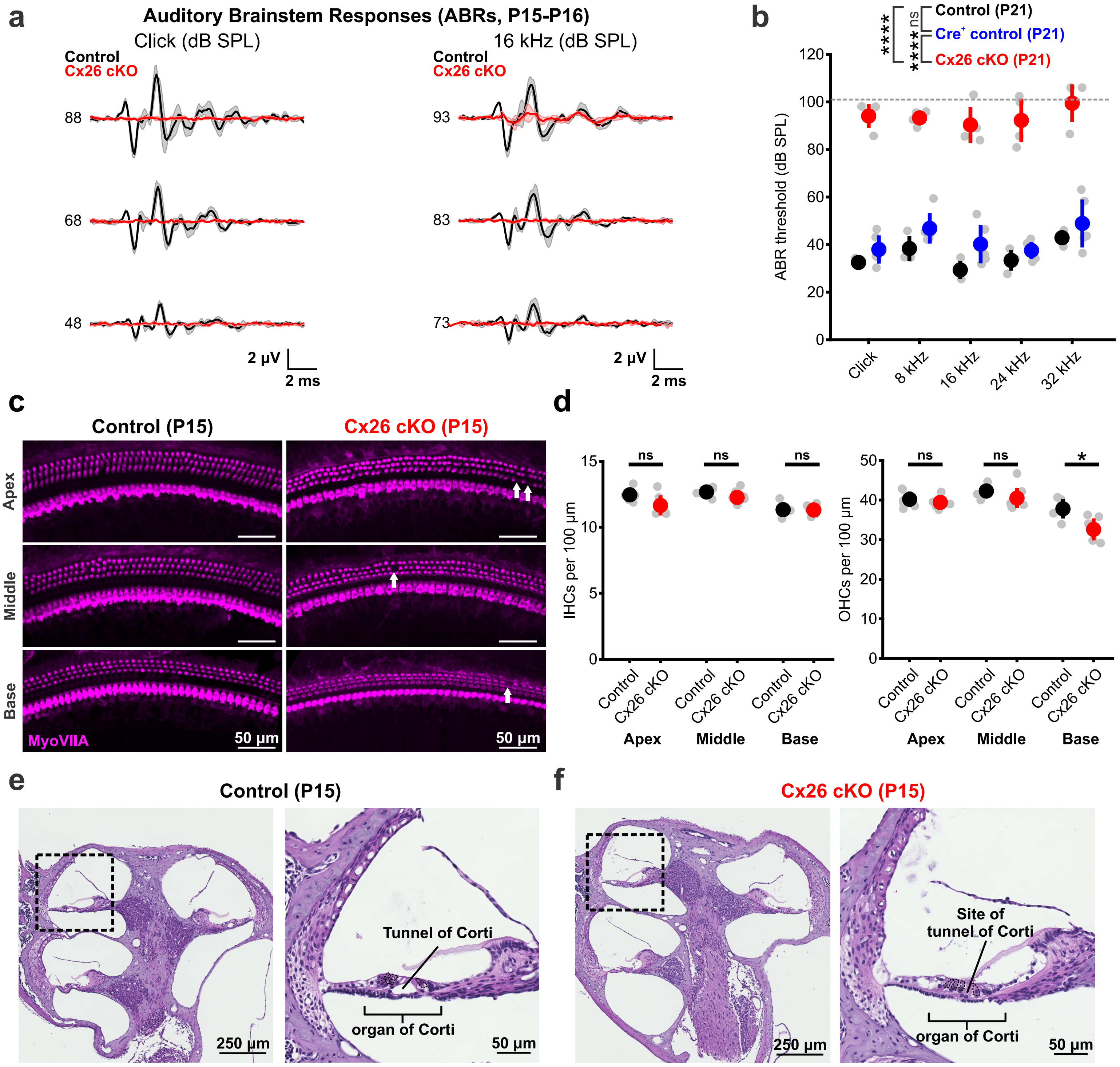


### Extended data Figure 8
